# Dichaete, a Sox2 homologue, prevents activation of cell death in multiple developmental contexts

**DOI:** 10.1101/2021.05.02.442335

**Authors:** Katherine Harding, Katerina Heath, Kristin White

## Abstract

Precisely regulated cell death plays a critical role in normal development and is controlled by the balance of pro-apoptotic and anti-apoptotic signals. In *Drosophila*, transcription of the clustered cell death activators *grim* and *reaper* is turned on in the developing nervous system to eliminate neural stem cells at the end of embryonic development. This transcription is activated by a pulse of the Hox gene *abdominal-A*. We show here that the Sox2 homologue *Dichaete* inhibits neural stem cell death when overexpressed, and loss of *Dichaete* promotes premature neural stem cell death. The anti-apoptotic activity of Dichaete opposes the pro-apoptotic factors *abdominal-A*, as well as the transcription factor *grainyhead*. The function of all three genes impinge on an enhancer that regulates the transcription of *grim* and *reaper*. Furthermore, we find that the balance between *abdominal-A* and *Dichaete* is likely to regulate the death of other cells during development, including cells in the developing midline, the developing hindgut, and in the early abdominal epidermis. Loss of *Dichaete* results in premature death in these tissues. This death can be rescued by the deletion of the enhancer region between *grim* and *reaper*. These data suggest that *Dichaete* functions to inhibit cell death activated by *abdominal*-*A* in multiple developmental contexts.

## INTRODUCTION

Programmed cell death is a critical element of normal development, with a notable role in shaping the nervous system. Apoptosis-deficient *Drosophila* mutants display reduced adult viability and a massively overgrown central nervous system due to disruption of the endogenous pattern of neural stem cell apoptosis (White et al, 1994; Tan et al, 2011). Similarly, a complete block in intrinsically activated apoptosis in mouse development, due to a triple knockout of the pro-apoptotic proteins BAX, BAK and BOK, results in lower than 2% survival to weaning and an expansion of multiple hippocampal layers during embryogenesis (Ke et al, 2018), suggesting that developmentally regulated apoptosis is required for organismal viability and normal brain development in vertebrates as well as *Drosophila*. As programmed cell death is an irreversible cellular behavior, the activation of this cell fate program must be under extremely tight regulatory control to avoid spurious activation. However, the organismal consequences of not activating cell death in the appropriate context are also dire, as this can lead to developmental defects or accumulation of transformed and malignant cells. Understanding how cell death is selectively and accurately activated in the appropriate context is therefore critical to our understanding of nervous system development.

We investigated the role of *Dichaete,* the *Drosophila* homologue of Sox2, in the regulation of neural stem cell death. Expression of mouse Sox2 is able to rescue phenotypes of *Dichaete* mutant embryos, indicating that these proteins are functionally conserved (Sánchez Soriano & Russell, 1998). Many lines of evidence have outlined critical roles for both *Dichaete* and Sox2 in neural development, but the consequences of loss of function of these proteins remain unclear. In early mammalian embryonic development, the transcription factor Sox2 is required for derivation of both the trophectoderm and inner cell mass in blastocysts (Avilion et al, 2003; Keramari et al, 2010; reviewed in Sarkar & Hochedlinger 2013). As Sox2-null embryos are therefore lethal prior to the generation of more differentiated tissues, hypomorphic and conditional knock-out alleles have been deployed to reveal functions for Sox2 in post-implantation development. During mammalian fetal development, Sox2 is primarily expressed in the neuroectoderm and gut endoderm, as well as the developing pharyngeal arches and germ cells (Sarkar & Hochedlinger 2013). Post-natal survival can be achieved in heterozygous mutant mice that express Sox2 protein at 25-30% of the wild type levels (Ferri et al, 2004). These adult mutant mice display pronounced neural phenotypes, including loss of cortical tissue associated with neurodegeneration and accumulation of intraneuronal aggregates (Ferri et al, 2004). The degenerating neurons in these Sox2-deficient brains display the hallmark characteristics of apoptotic cells, including condensed nuclei, hyperchromatic nuclear and intracellular staining, and membrane blebbing (Ferri et al, 2004). Similarly, neural-specific deletion of Sox2 during embryogenesis results in loss of neural stem cell populations by postnatal day 7, and this cell loss is associated with increased apoptosis throughout the dentate gyrus where post-embryonic neural stem cells are located (Favaro et al, 2009). Despite these observed links between Sox2 deficiency and apoptosis in neural stem cells, the role of cell death in driving Sox2 mutant phenotypes has remained elusive due to the difficulty in blocking apoptotic death in these animals.

The *Drosophila* homologue of Sox2, *Dichaete*, plays a similar role in neural development. It is highly expressed in the ventral neuroectoderm during embryonic development, and *Dichaete-*null alleles are embryonic lethal (Nambu & Nambu 1996; Sánchez Soriano & Russell, 1998). *Dichaete* mutant embryos display segmentation errors and significant defects in central nervous system development (Nambu & Nambu 1996; Sánchez Soriano & Russell, 1998). Previous mechanisms for these defects have been proposed, including a transcriptional role for *Dichaete* in specifying both midline glia and neural stem cell populations within the nervous system (Nambu & Nambu 1996; Sánchez Soriano & Russell, 1998; Ma et al, 2000; Zhao & Skeath 2002; Overton et al, 2002). However, as with mammalian phenotypes, the role of apoptotic cell death following *Dichaete* loss of function has not been investigated. Previous studies have suggested a role for *Dichaete* in regulating apoptosis: overexpression of *Dichaete* within the larval central nervous system resulted in neural stem cells that persist through a wave of developmental cell death (Maurange et al, 2008). However, the mechanism through which *Dichaete* may act to prevent activation of neural stem cell apoptosis remains unclear. We therefore investigated the connection between *Dichaete* and apoptosis, using the developmental cell death of *Drosophila* embryonic neural stem cells as a model system.

During embryonic development of the *Drosophila* nervous system, neural stem cells are generated in equal numbers within the thoracic and abdominal regions of the ventral nerve cord (VNC), resulting in approximately 30 neural stem cells per hemisegment. The majority of abdominal neural stem cells are eliminated by apoptosis towards the end of embryonic development, leaving a small number of abdominal neural stem cells that persist in the larval VNC (White et al, 1994; Harding & White 2018). Developmental apoptosis of neural stem cells requires the cell death genes *grim* and *reaper*, which are transcriptionally activated in the abdominal region of the embryo to promote apoptosis. Transcription of *grim* and *reaper* is controlled by an intergenic enhancer (enh1) that lies between the coding regions of these genes and that is removed in the *MM3* deletion background, resulting in ectopic abdominal neural stem cell survival in homozygous *MM3* mutants (Tan et al, 2011; Arya et al, 2015). The enh1 regulatory element is sensitive to activation by the pro-apoptotic factors *abdominal-A* (*abdA*) and *grainyhead* (*grh*), which are both necessary and sufficient for neural stem cell apoptosis (Prokop et al, 1998; Bello et al, 2003; Cenci & Gould, 2005; Maurange et al, 2008; Arya et al, 2015; Khandelwal et al, 2017). Grim and Reaper are RHG-domain containing proteins, which bind to the Drosophila Inhibitor of Apoptosis Protein 1 (DIAP1) resulting in increased DIAP1 turnover and release of bound initiator caspases, ultimately leading to cell death (Yang et al, 2000; Lisi et al, 2000; Suzuki et al, 2001; Wing et al, 2001; Ryoo et al, 2002; Yoo et al, 2002; Chai et al, 2003; Zachariou et al, 2003; Yokokura et al, 2004).

In this study, we took advantage of the stereotyped pattern of abdominal neural stem cell death in *Drosophila* to investigate the role of *Dichaete* in regulating apoptotic cell death. We find that overexpression of *Dichaete* is sufficient to block developmental cell death of abdominal neural stem cells, acting through the enh1 regulatory region. We also investigate the endogenous mechanism through which neural stem cell apoptosis is activated and find that expression of the pro-apoptotic Hox gene *abdA* distinguishes between surviving and doomed neural stem cells, rather than *Dichaete* levels. Finally, we show that multiple developmental defects of *Dichaete*-null embryos can be rescued by blocking ectopic cell death. Our findings suggest that both *Dichaete* and mammalian Sox2 may act as safeguards against activation of the apoptotic pathway in neural stem cells and in other tissues.

## METHODS

### Fly stocks

Flies were raised on standard cornmeal/yeast agar medium supplemented with live yeast. All crosses were performed at 25°C, embryos were collected on molasses plates supplemented with yeast paste. The following stocks were used in this study: *yw^67c23^, luc-RNAi* (BDSC #31603), *UAS-D* (BDSC #8861), *VAcht-GAL4* (BDSC #39220), *calx-GAL4/TM3* (BDSC #48160), *abdB-GAL4* (BDSC #49822), *UAS-GrhB/CyO* (BDSC #42227), *Grh.GFP* (BDSC #42272), *abdA-RNAi* (BDSC #35644). The *grim^C15E^ rpr^87^*, *Df(3L)MM2* and *Df(3L)MM3* stocks have been described previously (Tan et al, 2011). The following stocks were kind gifts: *UAS-p35* (Hay et al, 1994), *UAS-Nicd* (S. Artavanis-Tsakonas & Kazuya Hori), *dcr2, PCNA-GFP; wor-GAL4* (Arya et al, 2015; IK Hariharan), *UAS-Grim(III)* and *D^87^/TM3* (Nambu lab; Nambu & Nambu 1996; Wing et al, 1998) and *UAS-AbdA::HA* (Graba lab). The enh1(b+c)- GFP reporter construct incorporates the genomic sequence from 3L:18,359,842..18,362,329 (r6.39). The second chromosome *wor-GAL4, UAS-NLS-dsRed* line was generated previously in our lab (Arya et al, 2015). The third chromosome *wor-GAL4, UAS-NLS-dsRed* recombinant line was generated for this study from BDSC #56554 and #8547, and balanced over *TM6B,tub-GAL80* from BDSC #9490. The *sim-GAL4; UAS-NLS-dsRed* stock was generated for this study from BDSC #9150 (Scholz et al, 1997) and #8547. *D^87^* allele sequencing was done with the following primers: an amplicon containing the deletion was generated using *D87 US* 5’[TCTTACGCTATGGGCCAGGTAT]3’ with *D87 DS* 5’[TCGTCTGATTCCAGTCACAACA]3’, then sequenced with the internal primers *D seq 3* 5’[AAAGCGTGTTTCCATGTCCTTT]3’ and *D seq 6* 5’[TGGTTTGGAGGATTGGCTTACT]3’. Two independent recombinant *D^87^,MM3* lines were generated for this study and genotyped using the above *D^87^* primers and the following additional primer pairs:; *WH5’-* 5’[TCCAAGCGGCGACTGAGATG]3’ with *MM3 US2* 5’[TGTATGCTACCAGCGAGCAAAT]3’; *WH3’+* 5’[CCTCGATATACAGACCGATAAAAC]3’ with *MM3 DS2* 5’[TGACATTTGTCCAGGCTGACTT]3’.

### Embryo immunostaining

Primary antibodies used in this study: anti-AbdA (goat, 1:500) Santa Cruz dH17; anti-Antp (mouse, 1:100) DSHB 4C3; anti-axons (mouse, 1:5) DSHB BP102; anti-cDcp1 (rabbit, 1:100) Cell Signaling 9578; anti-cycB (mouse; 1:3) DSHB F2F4; anti-Dichaete (rabbit, 1:100) Nambu lab; anti-Dpn (rat, 1:150) Abcam ab195173; anti-dsRed (rabbit, 1:50) Clontech 632496; anti-Elav (mouse, 1:100) DSHB 9F8A9; anti-Engrailed (mouse, 1:2) DSHB 4D9; anti-GFP (chicken, 1:500) Abcam ab 13970; anti-GFP (rabbit, 1:250-1000) ThermoFisher A11122; anti-Grh (rabbit, 1:200) Bello lab; anti-Repo (mouse, 1:5) DSHB 8D12; anti-slit (mouse, 1:5) DSHB C555.6D. All secondary antibodies were obtained from ThermoFisher and used at 1:200 dilution in 1% nonfat milk. Briefly, embryos were dechorionated with 50% bleach, fixed with 4% formaldehyde in equal volume heptane and stored in 100% ethanol at -20°C if not used immediately. Embryos were rehydrated into PBS+0.1% Triton X-100 (PBST), blocked with 1% nonfat milk for 30min – 2h at room temperature prior to addition of primary antibodies. Primary incubations were performed for 2-5 days at 4°C in 1% nonfat milk, embryos were washed in PBST (>1h) and secondary incubations were performed for 2h at room temperature or overnight at 4°C in 1% nonfat milk. Embryos were washed with PBST (>1h), then rinsed with PBS and mounted with Fluoromount G mounting media. All embryos for quantitative fluorescent analysis measurements were processed in parallel.

### RNA fluorescent in situ hybridization

RNA-FISH was performed as described previously (Tan et al, 2011). 100ng DIG-labelled probe against *grim* or *reaper* transcript was used per sample (Sigma, 11-175-025-910). Probes were hybridized at 56°C overnight and subsequently detected by anti-DIG at 4°C overnight (1:1000; Roche 11-207-733-910) with secondary tyramide signal amplification for 2h at room temperature in the dark (AKOYA Biosciences, Cat NEL744001KT). Immunostaining was performed in conjunction with RNA-FISH: anti-GFP was incubated with samples at the same time as anti-DIG, and secondary antibody incubations were performed prior to TSA reactions.

### Imaging, quantification and statistical analysis

All imaging was done on a Nikon A1 confocal system with 1.0um between z-steps, scale bars are indicated on all images. All slides for a given experiment were imaged at the same confocal settings. All image analysis was performed using FIJI. Statistical analysis was performed using Graphpad Prism, all comparisons are unpaired parametric t-tests. Statistical levels of significance are indicated on figures as follows: * p<0.05, ** p<0.01, *** p<0.005, **** p<0.0001.

Quantification of cell numbers was performed by scrolling through all z-stacks and identifying Dpn+ nuclei within the region of interest (3 abdominal hemisegments, unless otherwise noted). Midline glia were analyzed in the same way and identified as Slit-holes within the midline cluster. Unless noted otherwise, fluorescent intensity measurements were performed on a single z-stack within the nuclear Dpn+ area, (except Fig 3 where PCNA-GFP was used to demarcate the cytoplasmic cell boundary) and normalized by subtracting background mean intensity of five areas of similar size within the same z-stack. For AbdA/Grh/D expression analysis in Fig 2, data were obtained from two technical replicates for each antibody staining. Values from each replicate of antibody staining were normalized to each other by centering on the mean expression level obtained from all cells at stage 13 in that replicate. Some values represent a single data point due to cell scarcity at later embryonic time points.

En stripe analysis in Fig 8 was performed on maximum projections using the “Plot Profile” function in FIJI along a segmented line, where each segment was perpendicular to the axis of the En stripe. All lines were 270um+/-5um long to avoid distortion of the signal along the X-axis. Hindgut length analysis in Fig 8 was performed on maximum projections using the line tool in FIJI, each measurement was taken from the anterior point of the posterior spiracles to the farthest anterior point of the hindgut En staining.

## RESULTS

### Dichaete inhibits neural stem cell apoptosis

Developmental cell death in *Drosophila* is regulated by the transcriptional activation of the RHG genes, including *grim* and *reaper*. To identify regulators of developmental cell death in the *Drosophila* embryo, we screened transcription factors and other DNA binding proteins for a role in regulating neural stem cell death (Arya et al, 2015). We identified *Dichaete* (*D*) as a candidate, consistent with the larval phenotype described in Maurange et al (2008). We overexpressed *Dichaete* with the neural stem cell driver *worniu-GAL4* (*wor-GAL4*) and found a significant increase in the number of cells expressing Deadpan (Dpn), a neural stem cell-specific transcription factor, in the abdominal region of the VNC at the end of embryogenesis (Fig 1A-B).

**Figure 1.**
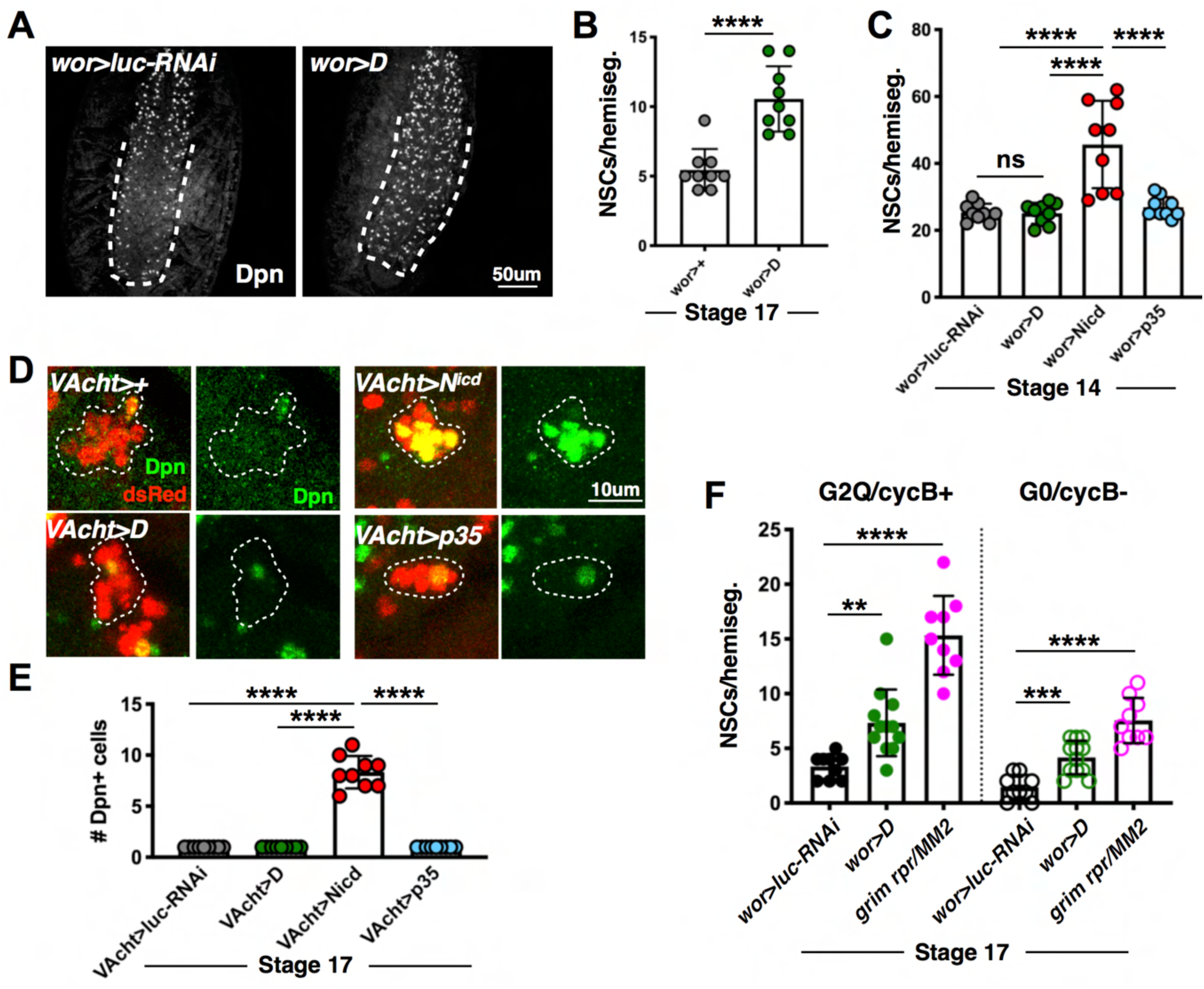
Dichaete is sufficient to inhibit abdominal neural stem cell apoptosis. **A)** Stage 17 embryos from *wor-GAL4,UAS-NLS-dsRed* (hereafter *wor>dsRed*) crossed to *luc-RNAi* or *UAS-*D and stained with anti-Dpn to visualize neural stem cells. Images are maximum projections through the VNC. **B)** Quantification of neural stem cell survival shown in **A**, n=3 hemisegments from 3 embryos per genotype. **C)** Quantification of abdominal neural stem cells in stage 14 embryos stained with anti-Dpn. n=3 hemisegments from 3 embryos per genotype. **D)** Abdominal NB3-5 dsRed+ progeny clusters in stage 17 embryos from *UAS-NLS-dsRed; VAcht-GAL4* flies crossed to indicated genotypes, stained with anti-Dpn. Images are maximum projections through the dsRed clusters. **E)** Quantification of Dpn+ cells shown in **D**, n=3 clusters from 3 embryos per genotype. **F)** Quantification of G2-quiescent (G2Q)/cycB+ and G0-quiescent (G0)/cycB- neural stem cell populations in stage 17 embryos. n=2 thoracic or 3 abdominal hemisegments from 3 embryos per genotype.

Multiple cellular processes could lead to the generation of ectopic neural stem cells in the VNC, including increased stem cell specification, inhibition of differentiation of neural progeny or the inhibition of abdominal neural stem cell death. To distinguish between ectopic neural stem cells generated through these processes, we assessed the effect of *Dichaete* on specification and differentiation. To address neural stem cell specification, we examined the number of Dpn+ neural stem cells at early stages of neurogenesis. While ectopic expression of a constitutively active form of Notch (*N^icd^*), known to promote neural stem cell identity (Bowman et al, 2008; Song & Lu, 2011; Zacharioudaki et al, 2012), significantly increased the number of Dpn+ cells observed at stage 14, we did not see any change in this early population when *Dichaete* was overexpressed or when cell death was inhibited with the caspase inhibitor p35 (Fig 1C). Next, we directly assessed whether *Dichaete* overexpression increased Dpn+ cell numbers by inhibiting the differentiation of neural progeny. We overexpressed *Dichaete* in a single neural stem cell lineage using the *VAcht-GAL4* driver (Lacin & Truman, 2016; Harding & White, 2019). The *Vacht-GAL4* driver is expressed in the neural stem cell NB3-5 and its progeny in each hemisegment. We have shown that NB3-5 is not eliminated by apoptosis (Harding & White, 2019), allowing us to evaluate the number of Dpn+ cells in this lineage at the end of embryogenesis. We did not see ectopic Dpn+ cells with overexpression of *Dichaete* or p35 within this lineage, whereas *N^icd^* was sufficient to generate additional Dpn+ cells (Fig 1D-E). Additionally, we tested whether ectopic neural stem cells in *Dichaete* overexpression embryos may be differentiated progeny that have failed to turn off the Dpn marker. However, we saw no co-expression of Dpn and either Repo (glial marker) or Elav (neuronal marker) in both control and *wor>D* embryos (Fig S1). Lastly, we recently described a novel terminal neural stem cell fate in the embryo that we have termed non-apoptotic loss (NAL; Harding & White 2019), which occurs specifically in G2 quiescent (G2Q) neural stem cells. We therefore tested whether *Dichaete* suppressed NAL, following our previous analysis. We found that *Dichaete* does not preferentially rescue the G2Q/cycB+ population that is subject to NAL (Fig 1F), indicating that ectopic neural stem cells in *wor>D* embryos do not result from inhibition of this novel fate. From these results, we conclude that the ectopic abdominal neural stem cells observed in embryos upon *Dichaete* overexpression are due to inhibition of apoptosis.

### AbdA levels, but not D or Grh, distinguish doomed and surviving neural stem cells

Activation of neural stem cell death has been shown to depend on the Hox gene *abdA* and the transcription factor *grh* (Prokop et al, 1998; Bello et al, 2003; Cenci & Gould 2005; Maurange et al, 2008; Arya et al, 2015; Khandelwal et al, 2017). We have now shown that *Dichaete* is sufficient to rescue embryonic neural stem cells from apoptosis, in addition to its previous characterization in larval development (Maurange et al, 2008). Integration of the roles of *abdA, grh* and *Dichaete* in cell death has been proposed in a previous model, whereby mutually exclusive Grh and D expression states signal the ability of a neural stem cell to undergo apoptosis: D+Grh- status is converted to D-Grh+ through a timing signal mediated by the temporal factor *castor*, and the D-Grh+ state is required for AbdA-mediated neural stem cell death (Maurange et al, 2008). Within this model, failure to repress *Dichaete* prevents downstream activation of RHG-mediated apoptosis, but the mechanism through which this occurred remained unclear.

To test this model directly, we took advantage of GAL4 driver lines generated by the Janelia Research Farms and characterized by Lacin & Truman (2016). We selected three GAL4 lines, *VAcht-GAL4*, *calx-GAL4* and *abdB-GAL4*, that are expressed in specific subsets of neural stem cells within each hemisegment of the VNC (Fig 2A), as described in Lacin & Truman (2016). These marked subsets include neural stem cells of known fate, including the surviving cell NB3-5 marked by *VAcht-GAL4* (described above), a mix of one surviving and four doomed cells marked by *calx-GAL4* and three doomed cells marked by *abdB-GAL4*. We used these drivers to express a nuclear dsRed fluorescent reporter and validated the neural stem cell assignments of these drivers by calculating the frequency with which an individual stem cell is observed per hemisegment at each stage of development. As the drivers have variable penetrance (Lacin & Truman, 2016), we normalized these frequencies to the maximum observed at any timepoint between embryonic stages 13-17 (plotted as red [doomed lineages] and green [survivor lineages] bars in Fig 2C-E). Our validation confirms that these driver lines are able to accurately identify individual neural stem cells of known fate *in vivo*, allowing us to use them to assess expression of cell death regulators in single cells over time.

**Figure 2.**
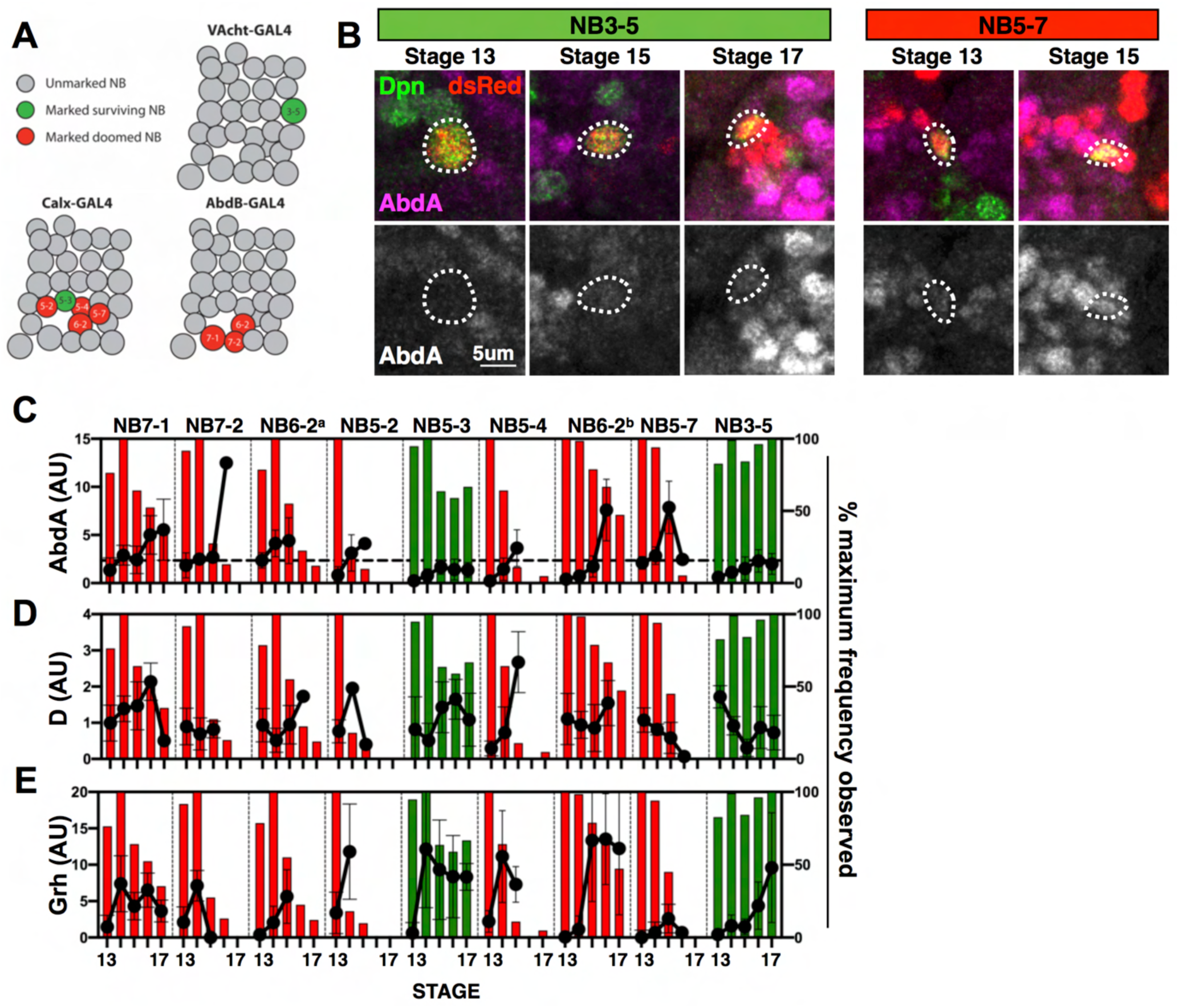
AbdA, not Grh or D, distinguishes doomed abdominal neural stem cells. **A)** Hemisegment map of neural stem cells with driver expression patterns indicated (from Lacin & Truman 2016; Harding & White 2019). **B)** Temporal dynamics of AbdA expression in the surviving neural stem cell NB3-5 (left; *VAcht>dsRed*) and the doomed neural stem cell NB5-7 (right; *calx>dsRed*). Neural stem cells are identified within reporter embryos by co-expression of Dpn and nuclear dsRed. Images are single confocal slices through the middle of the nuclear Dpn signal. **C-E)** Quantification of AbdA (**C**), D (**D**) and Grh (**E**) expression in neural stem cells during embryonic stages 13-17. Red (doomed cell) and green (surviving cell) bars represent survival frequency for each neural stem cell at a given embryonic stage (*VAcht>dsRed* st 13 n=10 embryos, st 14 n=17, st 15 n=20, st 16 n=14, st 17 n=19; *calx>dsRed* st 13 n=13 embryos, st 14 n=12, st 15 n=22, st 16 n=13, st 17 n=10; *abdB>dsRed* st 13 n=12 embryos, st 14 n=11, st 15 n=10, st 16 n=7, st 17 n=13). The dotted line in **C** indicates the highest level of AbdA measured in either surviving neural stem cell.

We used these drivers to quantify expression patterns of AbdA, D and Grh protein in individual neural stem cells from stages 13-17 of embryonic development. We drove a nuclear dsRed reporter and co-stained embryos for either AbdA, D or Grh, and Dpn to identify the neural stem cells. As an internal control, we correlated data points between replicate measurements of NB6-2, which is marked independently by *calx-GAL4* and *abdB-GAL4*. We found good correlation between these independent data sets (Fig S2), indicating that our strategy is able to accurately and reproducibly measure protein expression levels in individual neural stem cells.

We found that AbdA levels consistently rise to a higher peak in doomed cells just preceding the time of death, and this AbdA peak is not seen in the two surviving neural stem cells (Fig 2B-E). This suggests that AbdA levels alone are sufficient to distinguish between these two fates of neural stem cells. In contrast to the previously suggested model, we found no evidence for a consistent D+Grh- to D-Grh+ transition within individual doomed cells (Fig 2D-E). Grh was generally upregulated over time in all cells regardless of their fate, consistent with its documented role as a late member of the temporal series (Maurange et al, 2008; Kohwi & Doe, 2013; Harding & White 2018). D was more variably expressed over time, with no obvious correlation between levels of D and ultimate cell fate. To directly assess co-expression of Dichaete and Grh, we used a protein tagged GrhGFP allele and co-stained embryos for D, GFP and Dpn. The GrhGFP protein faithfully recapitulates endogenous Grh expression (Fig S3A). We find that surviving neural stem cells in the abdominal region of stage 17 embryos express both D and GrhGFP (Fig S3B), suggesting that there is no mutual exclusivity in expression of these transcription factors. With this new information in hand, we proceeded to investigate the mechanism through which *Dichaete* blocks neural stem cell apoptosis.

### Dichaete down-regulates transcription of cell death genes *grim* and *reaper*

As *Dichaete* is a transcription factor that has been shown to act variably as a transcriptional activator and repressor (Sarkar & Hochedlinger 2013), we hypothesized that *Dichaete* might act by regulating transcription of the cell death genes *grim* and *reaper* (*rpr*), which are required for neural stem cell apoptosis (White et al, 1994; Chen et al, 1996; Peterson et al, 2002; Tan et al, 2011). To assess *grim* and *rpr* expression, we used RNA fluorescent in situ hybridization (RNA-FISH; Tan et al, 2011; Arya et al, 2015; Arya et al, 2019). As we find that the Dpn protein epitope does not survive the RNA-FISH protocol, we used a PCNA-GFP reporter that marks proliferating cells as an alternative method to identify cell types in the developing nervous system: PCNA-GFP is expected to label both dividing neural stem cells and their immediate proliferative progeny, the ganglion mother cells (GMCs). We find that 60-64% of PCNA-GFP+ cells are Dpn+ at stage 14, and 46-48% at stage 15 in control and *wor>D* embryos, confirming that the PCNA-GFP reporter marks a similar population in both of these genetic conditions (Fig S4A). We found that overexpression of *Dichaete* decreased the level of both *grim* and *rpr* transcription in PCNA-GFP+ cells compared to control embryos (Fig 3A-D; Fig S4B-C). We confirmed that the lack of transcripts in *wor>D*-rescued PCNA-GFP+ cells is not due to rapid mRNA turnover in “undead” cells where apoptosis is blocked. Using the baculovirus caspase inhibitor p35 to block apoptosis, we found that *grim* transcript accumulates to significantly higher levels compared to *Dichaete* overexpression (Fig 3A-B), indicating that the *grim* transcript is not eliminated in undead cells. These results suggest that *Dichaete* inhibits apoptosis by preventing transcription of pro-apoptotic genes *grim* and *rpr*.

**Figure 3.**
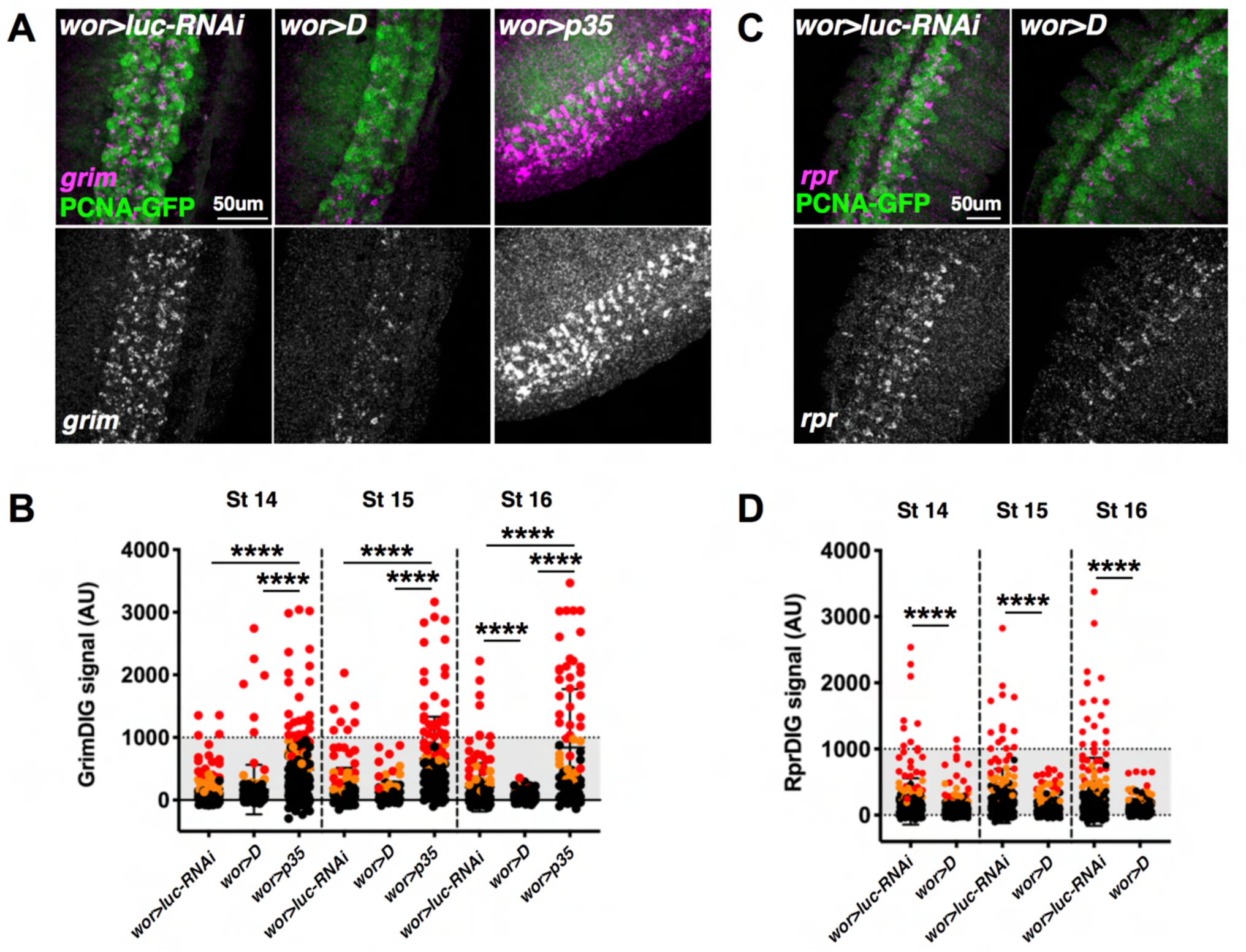
Dichaete downregulates *grim* and *rpr* transcription. **A)** Stage 15 embryos from *dcr2,PCNA-GFP;wor-GAL4* crossed to the indicated genotypes, stained for PCNA-GFP with RNA-FISH for *grim* transcript. Images are maximum projections through the VNC. **B)** Quantification of *grim* transcript intensity within PCNA-GFP+ cells shown in **A**. Values were measured in 3 abdominal segments of ≥2 embryos (*wor>luc-RNAi* st 14 n=227 cells, st 15 n=150, st 16 n=130; *wor>D* st 14 n=138, st 15 n=96, st 16 n=115; *wor>p35* st 14 n=203, st 15 n=128, st 16 n=86). Qualitative expression assignments are indicated by data point color: red – high expression, orange – low expression, black – no expression. **C)** Stage 14 embryos stained for PCNA-GFP with RNA-FISH for *rpr* transcript. Images are maximum projections through the VNC. **D)** Quantification of *rpr* transcript intensity shown in **C**, analysis as in **B**. (*wor>luc-RNAi* st 14 n=228 cells, st 15 n=194, st 16 n=187; *wor>D* st 14 n=323, st 15 n=192, st 16 n=158).

### Dichaete acts through the MM3 intergenic region to inhibit cell death

We have previously described a regulatory element (enhancer1; enh1) that is required for *grim* and *rpr* transcription and is found within the Neuroblast Regulatory Region (NBRR; Tan et al, 2011; Arya et al, 2015). To test whether *Dichaete* regulates *grim* and *rpr* transcription by altering activity of this regulatory element, we examined the effects of *Dichaete* overexpression and loss of function on enh1-GFP and enh1-dsRed reporters. These transgenic reporters of enh1 activity contain either 2.5kb (enh1-GFP) or 5kb (enh1-dsRed) of the cell death regulatory element, driving fluorescent protein expression. These reporters are specifically expressed in doomed neural stem cells in control embryos (Arya et al, 2015; Fig 4). We found that neural stem cells rescued by *Dichaete* overexpression are enh1-GFP-negative, suggesting that *Dichaete* acts upstream of enh1 (Fig 4A-B). In addition, we find that, while enh1-GFP signal accumulates in p35-rescued cells, co-expression of *Dichaete* and p35 significantly inhibits the expression of enh1-GFP in ectopic neural stem cells (Fig 4A-B). This result further indicates that *Dichaete* prevents the activation of the enh1-GFP transcriptional reporter of cell death.

**Figure 4.**
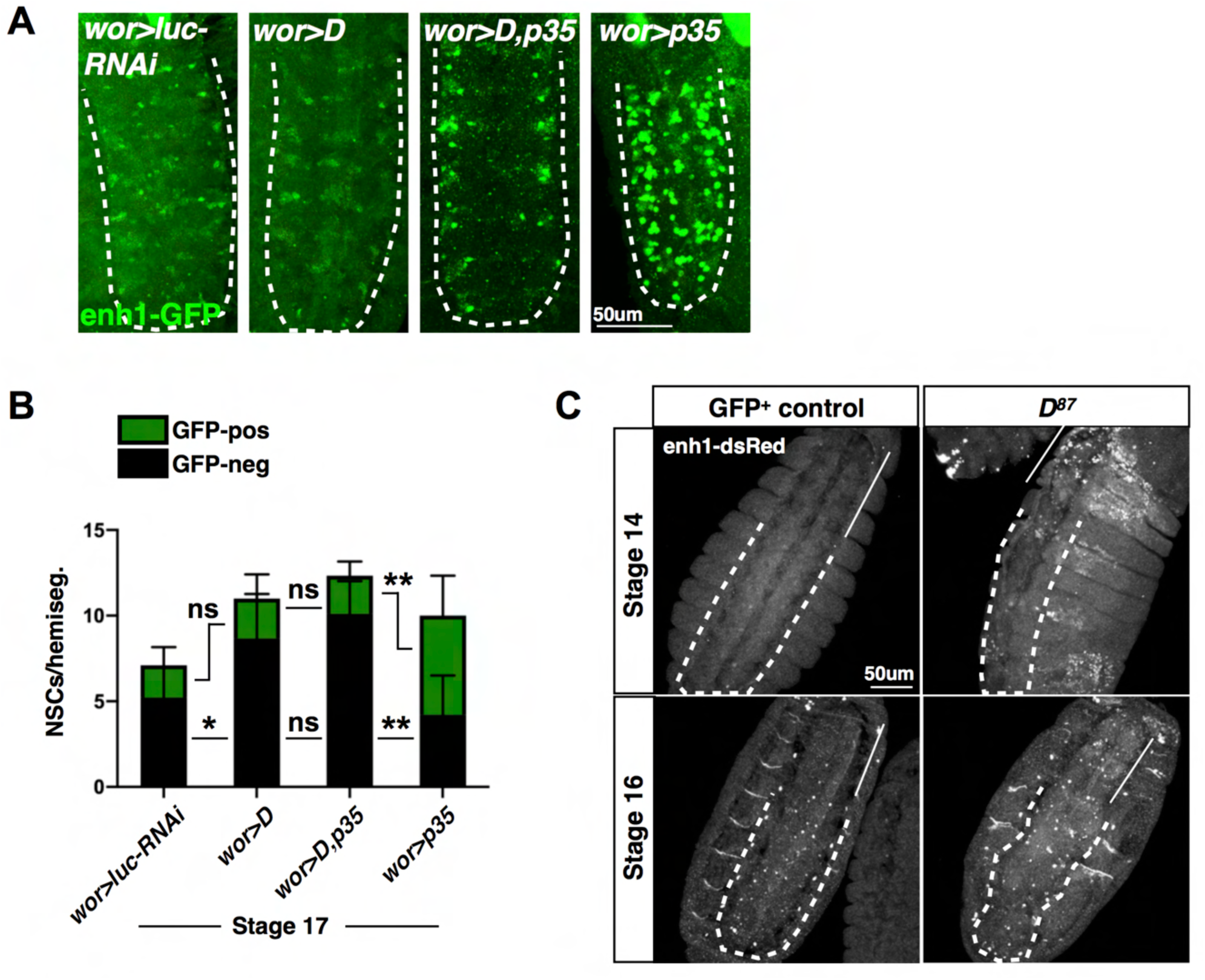
Dichaete regulates reporters of the enh1 cell death enhancer. **A)** Stage 17 embryos from *wor>dsRed;enh1(b+c)-GFP* crossed to the indicated genotypes, stained with anti-GFP to visualize enh1(b+c)-GFP reporter expression. The abdominal VNC is outlined with a white dashed line. Images are maximum projections through the VNC. **B)** Quantification of enh1(b+c)-GFP expression in Dpn+ abdominal neural stem cells at stage 17. n=3 hemisegments from 3 embryos per genotype. **C)** Stage 14 and 16 *enh1-dsRed;;D^87^/TM3,GFP* or *enh1-dsRed;;D^87^/D^87^* embryos stained with anti-dsRed. The thoracic region is indicated with a straight line and the abdominal region is outlined with a dashed line. Images are maximum projections.

To assess the endogenous relationship between *Dichaete* and activation of the cell death enhancer, we examined levels of Dichaete expression in enh1-GFP+ and enh1-GFP- neural stem cells in a wild type background. We also performed the same analysis with the pro-apoptotic factors AbdA and Grh. Consistent with our single neural stem cell analysis (Fig 2), we find that AbdA expression is significantly higher in the doomed enh1-GFP+ cells than the enh1-GFP- population at multiple time points (Fig S5A), while Grh and Dichaete generally show no significant difference in expression levels between these populations with the exception of stage 14 for Grh (Fig S5B-C). We note that while enh1-GFP expression is specific to doomed neural stem cells, not all dying neural stem cells express enh1-GFP, leading our measurements to be a likely underestimate of the expression differences of these factors between doomed and surviving neural stem cells.

We next examined the effect of *Dichaete* loss of function on enh1-dsRed reporter expression, using the *D^87^* allele generated by Nambu & Nambu (1996). We sequenced this allele and found that *D^87^* lacks 5’UTR and coding sequences of the *Dichaete* gene (deletion coordinates 3L:14,176,758..14,177,440 r6.38). In the *D^87^* mutant background, the enh1-dsRed reporter is strongly derepressed in the thorax of the embryo compared to sibling control embryos, and ectopic dsRed+ foci are observed throughout the thoracic and abdominal regions of the VNC (Fig 4C). Furthermore, at stage 14, enh1-dsRed expression is expanded outside of the nervous system to stripes in the epidermis. The thoracic domain of derepression coincides with the expression domain of Antp, which is normally co-expressed with Dichaete in early embryonic stages (Fig S6A). Dichaete is also co-expressed with AbdA at early embryonic stages in the neuroectoderm (Fig S6B). These results suggest that Dichaete normally functions to repress ectopic activation of the enh1 reporter by multiple Hox genes.

We find no evidence for ectopic upregulation or derepression of Grh or AbdA in the *D^87^* mutant embryos (Fig S7A-B), suggesting that the ectopic activation of the enh1-dsRed reporter does not occur through misregulation of these factors. Our results suggest that *Dichaete* is both necessary and sufficient for repression of the enh1 cell death regulatory element, while endogenous activation of the reporter is correlated with increased AbdA expression levels rather than downregulation of Dichaete.

### Midline glia cell death is *MM3*- and *Dichaete*-independent

The ability of *Dichaete* to regulate enh1 reporters does not rule out that it may also act at other genomic regions within the cell death locus to block apoptosis. To address this question, we tested whether *Dichaete* could block the apoptotic death of midline glia. Similar to neural stem cells, midline glia are generated in excess during embryogenesis and the majority are eliminated by cell death (Zhou et al, 1995; Dong & Jacobs, 1997). Within the embryonic nervous system, midline glia are the source of a secreted glycoprotein Slit (Rothberg et al, 1990). Apoptosis of midline glia is blocked by the *H99* deletion that removes the cell death genes *hid, grim* and *rpr* as well as the intervening sequences (White et al, 1994; Zhou et al, 1995; Dong & Jacobs, 1997). Deletion of *grim* and *rpr* alone in *grim rpr/MM2* mutants also inhibits midline glia death as detected by Slit staining (Fig 5A-B), confirming that some of this death is dependent on *grim* and *rpr*. However, it is unclear whether midline glia and neural stem cell apoptosis share regulatory elements. We therefore tested the effect of the intergenic *MM3* deletion on midline glia cell death. We found that *MM3* mutant embryos have wild type numbers of Slit+ midline glia, indicating that midline glia cell death is *MM3*-independent (Fig 5A-B). We next tested whether *Dichaete* misexpression can block this form of *MM3-*independent cell death using the *sim- GAL4* driver combined with nuclear dsRed (Scholz et al, 1997; Sánchez Soriano & Russell, 1998). We find that *Dichaete* overexpression does not block midline glia death, whereas blocking death with *sim>p35* resulted in significant rescue of midline glia (Fig 5C-D). *Dichaete* therefore cannot block this *MM3-*independent apoptosis. From these findings and those above, we conclude that *Dichaete* inhibits cell death by preventing activation of the enh1 regulatory region, and may not block cell death by acting through additional regions of the cell death locus.

**Figure 5.**
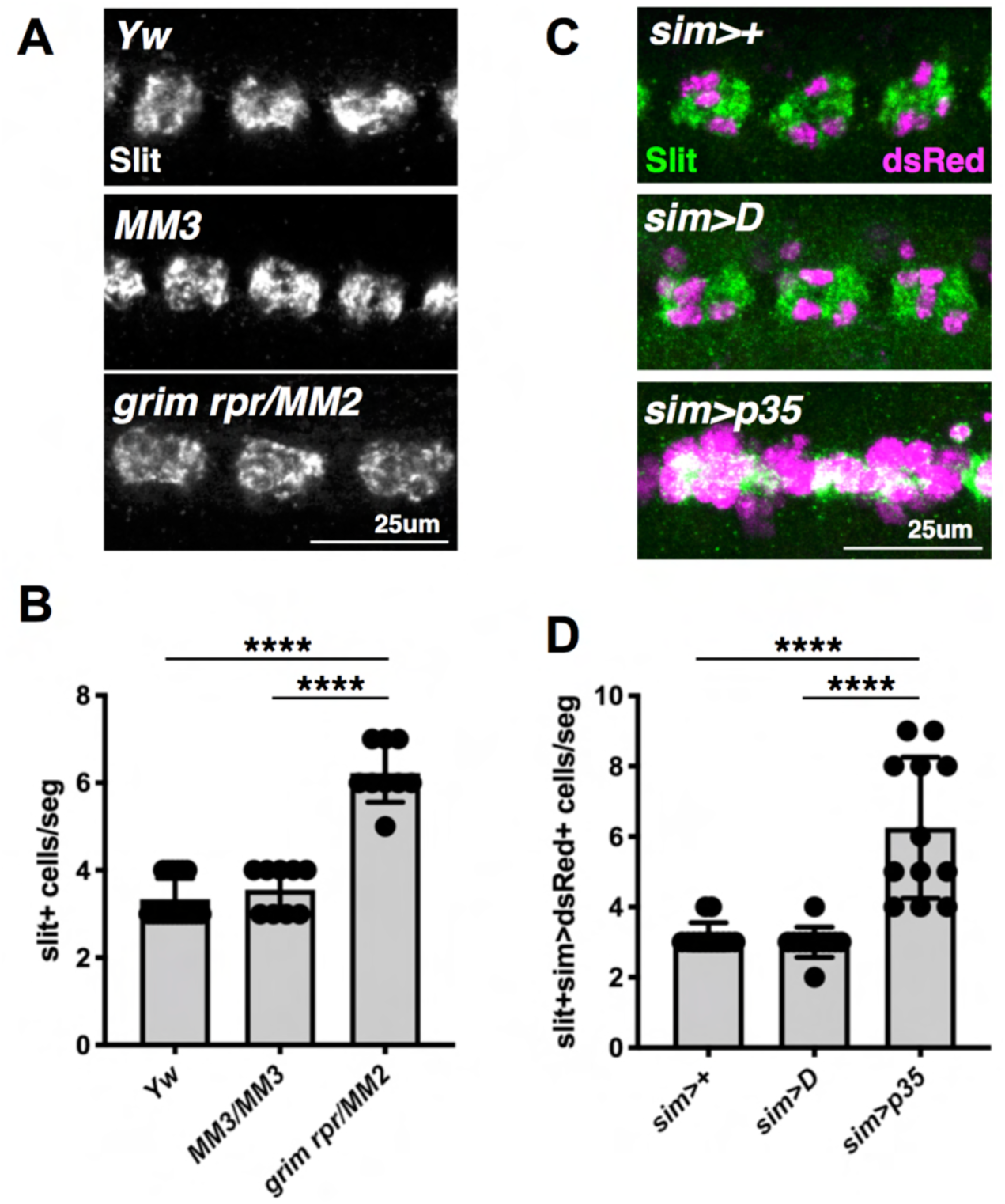
Midline glia death is *MM3-* and *Dichaete-*independent. **A)** Stage 17 embryos stained with anti-Slit to visualize midline glia clusters. Images are maximum projections through the Slit signal. **B)** Quantification of Slit+ cells within each midline cluster, shown in **A**. n=3 clusters from 6 (*Yw*)or 3 (*grim rpr/MM2* and *MM3*) embryos. **C)** Stage 17 embryos from *sim-GAL4;UAS-NLS-dsRed* crossed to the indicated genotypes, stained with anti-Slit. Images are maximum projections through the Slit signal. **D)** Quantification of Slit+dsRed+ nuclei per midline cluster, shown in **C**. n=3 clusters from 4 embryos per genotype.

### Dichaete acts downstream of AbdA and Grh to block cell death

As AbdA and Grh have been shown previously to regulate the enh1 element (Arya et al, 2015; Khandelwal et al, 2017), we wondered how the role of *Dichaete* in blocking neural stem cell death intersected with these pro-apoptotic regulators. We first tested the effect of *Dichaete* overexpression on levels of AbdA and Grh. We found that *Dichaete* overexpression significantly downregulated expression of both AbdA and Grh (Fig S8A-B), suggesting that *Dichaete* inhibits neural stem cell death by turning down pro-death signalling through AbdA and Grh. We note that the decrease in AbdA levels in *wor>D* embryos is still significantly higher than those seen with *wor>abdA-RNAi*, however both genotypes have similar neural stem cell survival at stage 17 (Fig S8C). To directly test the functional relationship between AbdA/Grh and Dichaete, we co-expressed *Dichaete* and either *abdA* or *Grh* and assessed neural stem cell survival. Interestingly, we found that *Dichaete* was able to rescue neural stem cell apoptosis even in the context of *abdA* or *Grh* overexpression (Fig 6A-B). Both *abdA* and *Grh* are sufficient to kill thoracic neural stem cells with the *wor-GAL4* driver (Fig S9), indicating that they ectopically activate cell death in these conditions. We also find that *abdA* and *Grh* are not sufficient to kill midline glia (Fig S10A-B), nor does RNAi knockdown of *abdA* lead to ectopic midline glia (Fig S10C-D). Taken together with the finding that the *MM3* region is not required for midline death, these data suggest that, as with *Dichaete,* the mechanism through which *abdA* and *grh* activate cell death requires sequences within the *MM3* region. Together, our findings suggest that *Dichaete* is able to block cell death through two mechanisms: downregulation of pro-apoptotic factors AbdA and Grh, and inhibition of AbdA/Grh activity within the *MM3* region (Fig 6C).

**Figure 6.**
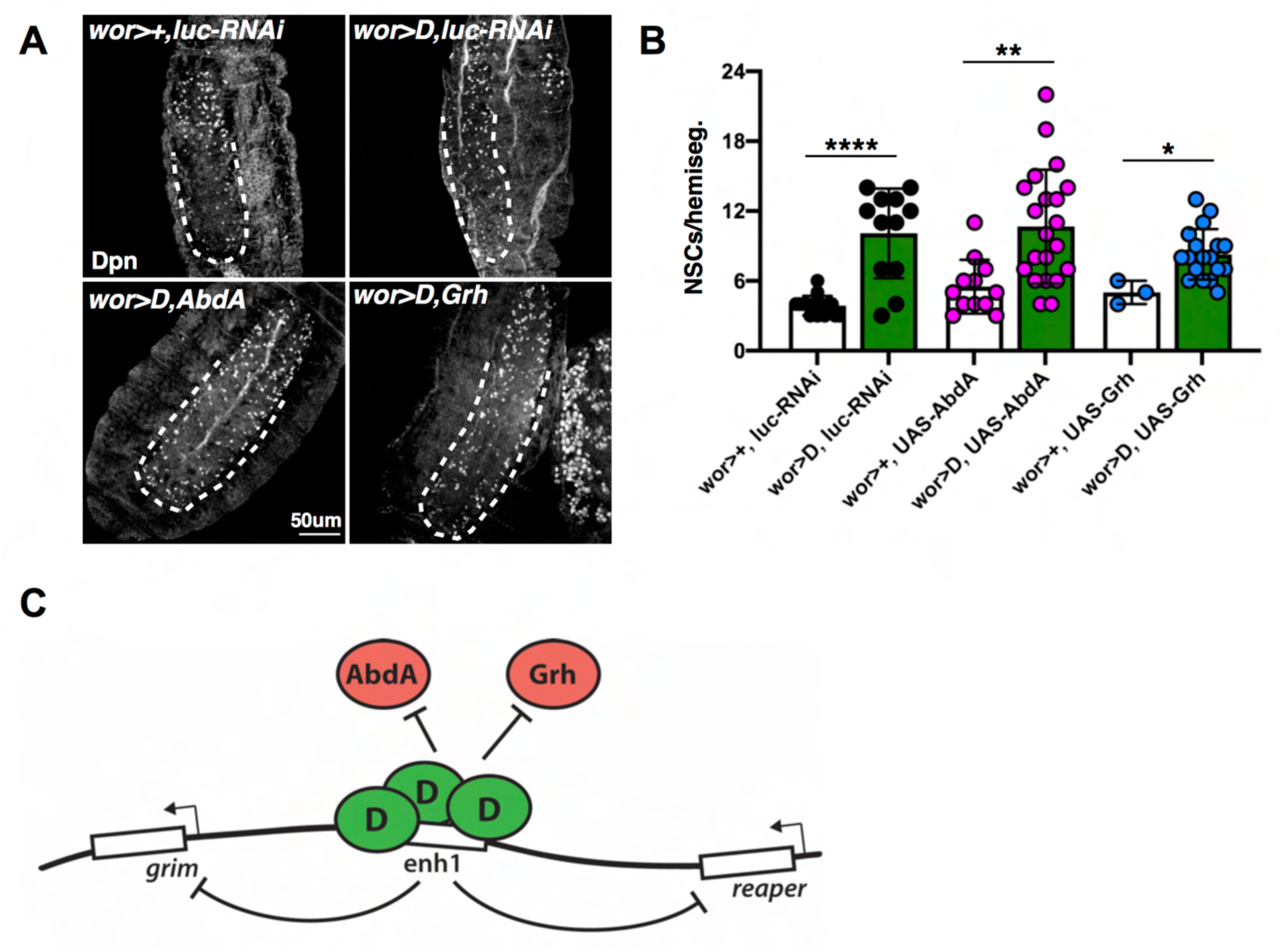
Dichaete blocks cell death downstream of pro-apoptotic factors AbdA and Grh. **A)** Stage 17 embryos from *wor>dsRed* or *UAS-D;wor>dsRed/TM6B,tubGAL80* crossed to *luc-RNAi, UAS-AbdA::HA* or *UAS-Grh*, stained with anti-Dpn to visualize neural stem cells. Images are maximum projections through the VNC. **B)** Quantification of neural stem cell survival in abdominal hemisegments as shown in **A**, n=3 abdominal hemisegments of ≥1 embryos. **C)** Model for role of Dichaete in neural stem cell apoptosis. Dichaete overexpression inhibits cell death at least in part by preventing activation of the cell death enhancer, enh1, by AbdA and Grh. Inhibition of enh1 then blocks downstream transcriptional activation of the cell death genes *grim* and *rpr*.

### Dichaete mutant phenotypes are rescued by blocking cell death

Thus far, our findings have pointed to a model in which *Dichaete* acts upstream of the enh1 regulatory region to inhibit neural stem cell death. To test whether *Dichaete* also functions downstream of this regulatory element, we generated recombinant alleles of *D^87^* and *MM3* to examine neural stem cell survival in this background. Consistent with previous reports (Overton et al, 2002; Zhao & Skeath, 2002), we find that *D^87^* mutant embryos display a loss of abdominal neural stem cells at stage 14 (Fig S11A), however we find that these mutants show wild type neural stem cell numbers at stage 17 (Fig 7A-B). Two independent recombinant *D^87^*,*MM3* alleles completely recapitulated the *MM3* neural stem cell survival phenotype (Fig 7A-B), indicating that *Dichaete* is not required to block cell death downstream of *MM3*. Interestingly, the *D^87^*,*MM3* embryos also rescue the reduction in neural stem cell numbers observed in *D^87^* mutant embryos at stage 14 (Fig S11A), suggesting that the loss of neural stem cells in early *D^87^* embryos is due to ectopic activation of cell death. We note that these defects are restricted to the AbdA domain, whereas numbers of neural stem cells in the thorax of stage 14 *D^87^* embryos are normal (Fig S11B).

**Figure 7.**
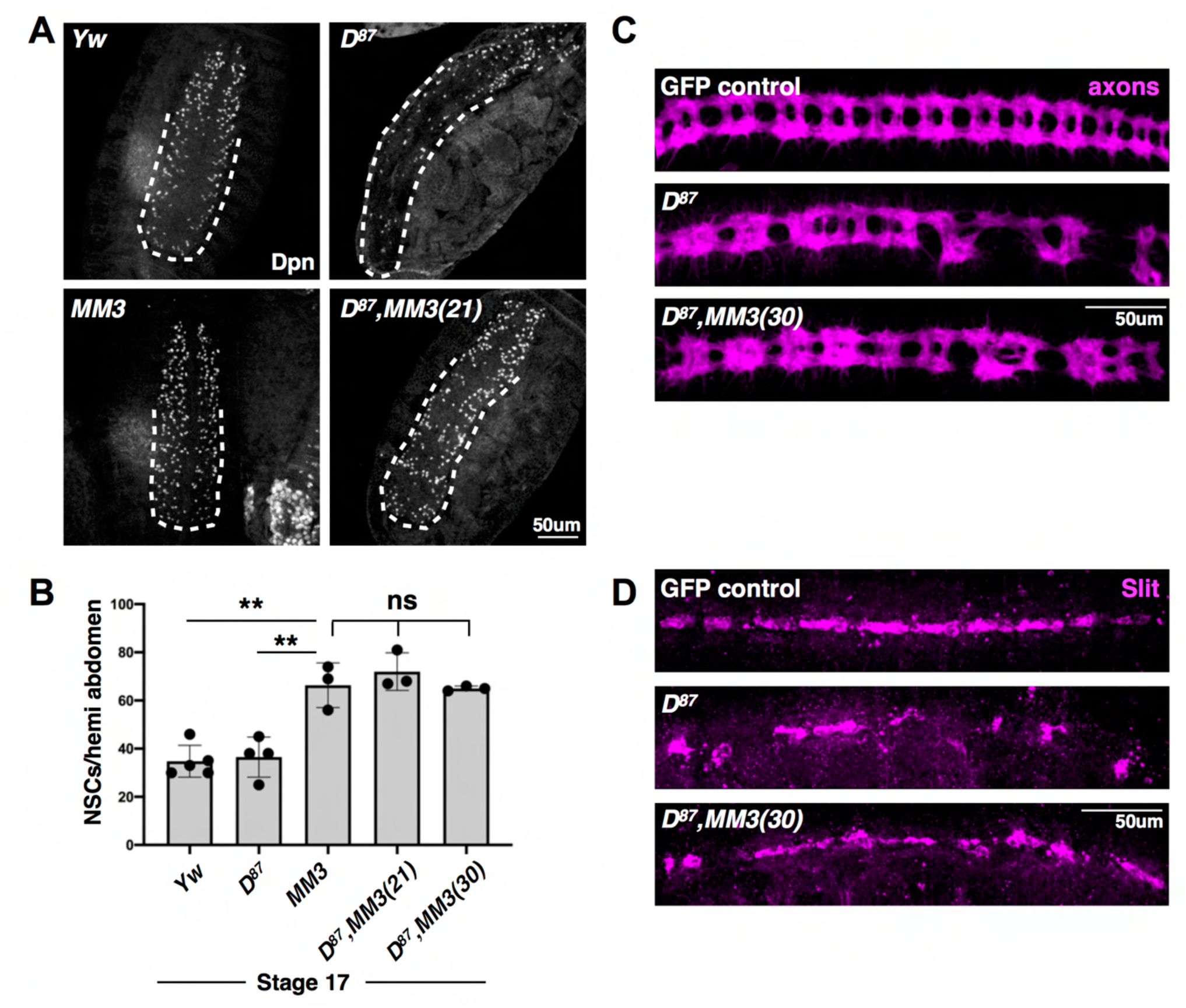
Deletion of cell death regulatory region rescues *D^87^* central nervous system defects. **A)** Stage 17 embryos stained with anti-Dpn to visualize neural stem cells. Images are maximum projections through the VNC. **B)** Quantification of abdominal neural stem cell survival as shown in **A**. Dpn+ cells were counted in hemi-abdomens as the abdominal segmentation defects in *D^87^* backgrounds preclude accurate hemisegment analysis at this stage. n≥3 embryos per genotype. **C)** Stage 16 embryos stained with anti-axons to visualize axonal tracts. Images are maximum projections through the axons signal. **D)** Stage 14 embryos stained with anti-Slit to visualize midline glia clusters. Images are maximum projections through the Slit signal.

In addition to neural stem cell phenotypes, *D^87^* mutants display significant axonal disorganization, including narrowing of longitudinal connectives between segments and fusion of commissural axons across the midline (Nambu & Nambu 1996; Sánchez Soriano & Russell, 1998). We therefore tested whether axonal morphology is rescued in *D^87^,MM3* embryos by staining for axon tracts with the monoclonal BP102 antibody (Elkins et al, 1990; Nambu & Nambu 1996; Sánchez Soriano & Russell, 1998). We find that *D^87^,MM3* recombinant embryos have improved organization of both longitudinal and commissural axon tracts relative to *D^87^* mutants alone, but do not fully recapitulate wild type organization (Fig 7C). As *Dichaete* expression in midline glia alone is sufficient to rescue axonal defects of *Dichaete* null embryos (Sánchez Soriano & Russell, 1998), we examined midline glia populations in *D^87^*,*MM3* embryos and found rescue of Slit+ cells along the midline relative to the *D^87^* mutant alone (Fig 7D). Again we find that the *D^87^*,*MM3* background does not completely recapitulate wild type organization of midline glia, consistent with our observation of improved but not completely rescued axonal morphology.

Our previous data suggests that Dichaete is required to inhibit AbdA-activated death. We observe that AbdA is expressed along the midline of wild type stage 11 embryos (Fig S12A), suggesting that loss of *Dichaete* leads to ectopic, AbdA-mediated death of midline cells. We did not observe AbdA and Slit co-expression at stage 15 (Fig S12B), consistent with our observation that normal midline glia cell death is AbdA-independent. A previous report has shown that *Dichaete* directly regulates *slit* expression and contributes to expression of *sim*, which is required for the formation of midline glia (Nambu et al, 1991; Ma et al, 2000). Our results indicate that the midline glia and Slit expression can be rescued in *D^87^* mutants by blocking cell death, suggesting that Slit expression can still be maintained downstream of *Dichaete.* This could be mediated by factors such as *sim* or *ventral veins lacking* (*vvl*, also referred to as *drifter)*, which cooperate with *Dichaete* to specify the midline glia (Ma et al, 2000). Together with the findings above, these results suggest that the primary neural defect in *D^87^* mutant embryos is ectopic activation of cell death, resulting in loss of neural stem cells and midline signaling.

As *D^87^* mutants display defects in segmentation (Nambu & Nambu 1996; Sánchez Soriano & Russell, 1998), we tested whether loss of *Dichaete* led to ectopic activation of cell death in stage 10/11 embryos by assessing levels of active, cleaved Dcp1 caspase (cDcp1). We found that *D^87^* mutant embryos at this stage display ectopic cDcp1 staining compared to control embryos, indicating that cell death is precociously activated in the absence of *Dichaete* (Fig S13). Notably, ectopic cDcp1 staining is largely absent from the thoracic segments of *D^87^* mutants but present in the head and abdominal regions. To determine if the segmentation defects are due to cell death, we examined segmentation patterns in *D^87^* and *D^87^*,*MM3* mutants by staining for Engrailed (En), which is expressed at the posterior border of embryonic segments. We measured En intensity along 9 stripes from anterior to posterior and found a periodic pattern of staining with regular spacing in control embryos using this method (Fig 8A-B). In contrast, the En stripes in *D^87^* mutants are disrupted and irregular, including missing segments and stripes that have fused together as has been described previously (Fig 8A-B; Nambu & Nambu 1996). We found that the striped En pattern is largely rescued in the *D^87^*,*MM3* background relative to *D^87^* mutants, restoring the 9 stripe periodicity of En staining in our quantification (Fig 8A-B). This suggests that the segmentation errors observed upon loss of *Dichaete* are due to ectopic cell death, activated by regulatory elements deleted in *MM3*.

**Figure 8.**
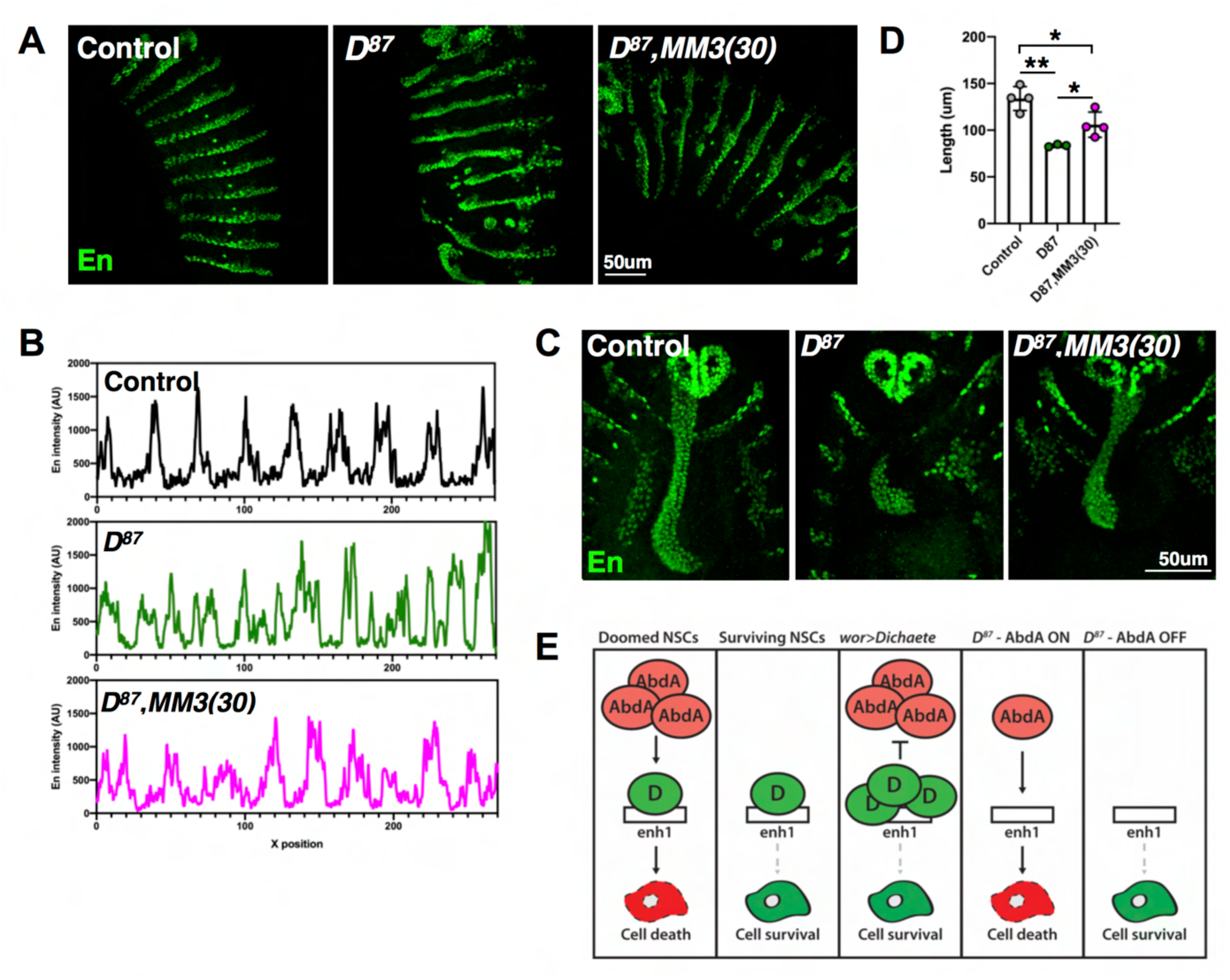
Multiple tissue defects of *D^87^* are rescued by blocking *MM3-*dependent cell death. **A)** Stage 13 embryos stained with anti-En to visualize posterior segment borders. Images are maximum projections. **B)** Quantification of En fluorescent intensity along 9 stripes as seen in **A**. Traces are average values from 4 embryos per genotype, left-right X-position is anterior-posterior. **C)** Stage 14 embryos stained with anti-En to visualize the developing hindgut. Images are maximum projections through the En signal. The loss of En staining in the hindgut of *D^87^* embryos as seen here has also been noted previously (Sanchez-Soriano & Russell, 2000). **D)** Quantification of hindgut length as shown in **C**, measured from the anterior point of the En+ posterior spiracles to the farthest anterior point of En signal. n=3-4 embryos per genotype. **E)** Model figure for the role of Dichaete in regulating activation of cell death through enh1. The apoptotic death of doomed neural stem cells is triggered by expression of AbdA, whereas surviving neural stem cells do not express AbdA and are therefore retained during post-embryonic development. The presence of ectopic D in *wor>D* embryos prevents the normal activation of enh1 by high levels of AbdA in doomed neural stem cells, leading to an increase in neural stem cell survival. Loss of Dichaete function allows precocious activation of enh1 in tissues where AbdA is expressed, whereas loss of Dichaete does not result in significant cell death in the absence of AbdA expression.

In addition to segmentation and neural phenotypes, *D^87^* mutants also exhibit hindgut defects (Sánchez Soriano & Russell, 2000). *Dichaete* is expressed in the primordium of this tissue and is required for proper development of the hindgut (Sánchez Soriano & Russell, 2000). However, the mechanism through which loss of *Dichaete* prevents normal development of the hindgut is unclear. As noted above, the defects of *D^87^* embryos are prominent within the domain of AbdA expression. We stained wild type embryos for Dichaete and AbdA, and found that they are co-expressed in the hindgut primordium at stage 10 (Fig S14). We therefore tested whether hindgut defects of *D^87^* embryos are rescued in the *D^87^*,*MM3* background, by again staining for En which marks the posterior gut (Sánchez Soriano & Russell, 2000; Takashima & Murakami, 2001). We found that *D^87^*,*MM3* embryos exhibit significant rescue of hindgut tissues relative to *D^87^* embryos (Fig 8C-D). Together these results lead us to propose that *Dichaete* loss of function in multiple tissues leads to the ectopic activation of cell death, resulting in posterior segmentation defects and tissue loss.

## DISCUSSION

In this work, we have investigated the function of the transcription factor *Dichaete* in regulating cell death in the *Drosophila* embryo. We have found that *Dichaete* is sufficient to inhibit developmental cell death of neural stem cells, and that *Dichaete* overexpression is associated with transcriptional downregulation of the cell death genes *grim* and *rpr*. The results obtained with our enhancer reporter and additional genetic data indicate that *Dichaete* blocks cell death by inhibiting activation of the enh1 regulatory region. Consistent with this model, we find that *Dichaete* downregulates expression of both AbdA and Grh, pro-apoptotic factors that are known to regulate enh1. Interestingly, we find that *Dichaete* can also block cell death downstream of AbdA and Grh, suggesting multiple mechanisms through which high levels of *Dichaete* interfere with activation of the apoptotic pathway. We also examined loss of function phenotypes of *Dichaete* and have shown that a primary defect in multiple tissues of *D^87^* mutant embryos is the ectopic activation of cell death, particularly in tissues that express AbdA. Blocking apoptosis with the intergenic *MM3* deletion is sufficient to rescue neural, segmentation and hindgut phenotypes observed in *D^87^* mutant embryos. This work describes a novel role for *Dichaete* in inhibiting apoptosis by regulating the enh1 cell death regulatory region and demonstrates that ectopic cell death contributes to a range of phenotypes observed upon loss of *Dichaete* function.

### Model for Dichaete-mediated protection against AbdA

Our work has identified a role for *Dichaete* in preventing ectopic activation of the cell death pathway. Our analysis of Dichaete and AbdA protein expression patterns in early embryos indicates that Dichaete is expressed constitutively throughout the epidermis prior to the onset of AbdA protein expression (Fig S6). Later, these proteins are expressed in distinct but overlapping domains in the embryo. The earliest embryonic defects arise in *D^87^* mutants around stage 5, where the expression pattern of the pair-rule gene *fushi tarazu* (*ftz*) is disrupted (Nambu & Nambu, 1996). The location of this disruption overlaps with the domain of early *abdA* transcript expression (Berkeley Drosophila Genome Project; Tomancak et al, 2002), suggesting a correlation between *abdA* expression and morphological defects upon loss of *Dichaete*.

We suggest a model in which the balance of anti-apoptotic signaling from *Dichaete* and pro-apoptotic signaling from *abdA* are integrated at enh1 (Fig 8E). Endogenous activation of abdominal neural stem cell death is associated with a pulse of high AbdA protein expression, which does not occur in surviving neural stem cells. We observe that *Dichaete* overexpression is sufficient to block *abdA*- mediated killing of neural stem cells (Fig 6). This suggests that overexpression recapitulates the high levels of Dichaete normally found in the neuroectoderm of early embryos, where an early wave of AbdA is expressed (Fig S6). We also show that *Dichaete* is not essential for all neural stem cell survival, as when we examined the effect of *Dichaete* loss of function, we did not observe ectopic death of thoracic neural stem cells or abdominal neural stem cell at the end of embryogenesis (Fig 7; Fig S11). As described above, in the two surviving neural stem cells that we are able to identify, we do not observe high AbdA expression. Thus, the loss of *Dichaete* in these cells does not lead to their ectopic death, as they do not express AbdA. We have now shown that morphological defects in multiple tissues are primarily the result of ectopic cell death, indicating that *Dichaete* normally functions to specifically protect against cell death activation associated with early expression of *abdA*. Our work indicates a context-specific requirement for *Dichaete* function in preventing cell death, associated with protection against high *abdA* levels.

### Regulation of cell death genes by enh1

Our work here and in previous studies has pointed to the enh1 regulatory region as a hub for cell death signal integration (Tan et al, 2011; Arya et al, 2015; Arya et al, 2019), leading to timely and accurate activation of cell death gene expression in neural stem cells. Additional work from others has determined that sequences within the enh1 region can be directly bound by AbdA and Grh, leading to the formation of a multi-protein tetracomplex on DNA *in vitro* (Khandelwal et al, 2017). While not investigated in this study, there is strong evidence for direct binding of Dichaete to the enh1 regulatory region as well, as determined by ChIP and DamID (modENCODE, 2010; Negrè et al, 2011; Aleksic et al, 2013). Taken together, this evidence suggests that physical binding of Dichaete to enh1 may prevent its recognition by the pro-apoptotic factors AbdA and Grh. This model is consistent with our finding that *Dichaete* overexpression is sufficient to block AbdA- or Grh-mediated killing (Fig 6). Similarly, we observe that loss of *Dichaete* leads to derepression of the enh1-dsRed reporter (Fig 4C), indicating that *Dichaete* is normally required to suppress activity of this enhancer. We can imagine two mechanisms through which Dichaete could function to block AbdA activity at enh1: Dichaete binding itself may physically prohibit subsequent AbdA binding, or Dichaete may recruit additional factors that serve to create a non-permissive environment around enh1.

In this work, we also find that *Dichaete* overexpression is sufficient to downregulate transcription of the cell death genes *grim* and *rpr* (Fig 3). We have previously shown that the *MM3* region containing enh1 is required for expression of *grim* and *rpr* in neural stem cells (Tan et al, 2011). However, it remains unclear how the regulatory signals at the enh1 hub are relayed to the promoters of *grim* and *rpr*. We have recently shown that multiple cohesin complex members are required for neural stem cell death (Arya et al, 2019), implicating long-range chromatin interactions in the activation of the cell death genes. Interestingly, recent work detected tissue-specific DNase hypersensitivity peaks at enh1, *grim* and *rpr* in *worniu-GAL4*-expressing cells at 4-6h, many hours before activation of neural stem cell death (Reddington et al, 2020). Consistent with recent datasets obtained from single cell ATAC- and RNA-seq (Bonn et al, 2012; Cusanovich et al, 2018; Reddington et al, 2020), the DNase hypersensitivity peaks observed within the cell death locus prior to death indicates that accessibility of the enhancer and promoters is not sufficient for gene expression, and that opening of the chromatin region occurs prior to gene activation. It will be of great interest in the future to examine if chromatin states of enh1 and the *grim/rpr* promoters are coupled, potentially through their physical interactions *in vivo*, and what mechanisms mediate transmission of regulatory input at enh1 to the gene promoters.

### Differential integration of Hox signaling

As noted above, we observed strong de-repression of the enh1-dsRed reporter in the absence of *Dichaete* function (Fig 4C). This derepression was observed in both the thoracic and abdominal regions of the embryo, however we did not observe a concomitant activation of ectopic cell death in the thorax: thoracic neural stem cell counts are normal at stage 14 in *D^87^* embryos and thoracic segmentation defects were less pronounced than those in the posterior. Our observations are entirely consistent with the initial phenotypic description of the *D^87^* allele by Nambu & Nambu (1996), which found no effect on thoracic denticle belts in *D^87^* embryos and noted that segmentation defects were largely restricted to the abdominal region. The observed discrepancy in coupling of enh1-dsRed reporter expression to activation of the cell death genes could reflect endogenous differences in the function of the two Hox genes expressed in these domains, Antp and AbdA.

While Hox genes have been studied extensively for decades, very little is known about the mechanism through which this family of transcriptional regulators interfaces with the transcriptional machinery. The few existing studies have suggested that Hox proteins variously interact with RNA Polymerase II-associated factors, including Mediator complex members Med13 and Med19 (Boube et al, 2000; Boube et al, 2014), TAF3 (TATA-activating factor 3; Prince et al, 2008) and the transcription pausing factor M1BP (Zouaz et al, 2017). In a direct comparison of Med19 binding to multiple Hox proteins, the Hox domain of Antp protein was sufficient for strong binding to Med19, whereas a weaker interaction was observed for AbdA (Boube et al, 2014). Mining of protein interaction databases has also suggested that Antp and AbdA interact with distinct general transcription factors (reviewed in Rezsohazy et al, 2015), potentially arguing for functional differences in their ability to couple enhancer signaling to transcriptional activation of downstream target genes such as *grim* and *reaper*. However, to date there has been no systematic analysis of differential Hox protein interactions with the transcriptional machinery, and the identity of factors that may contribute to selectivity of the effects of Antp and AbdA are unknown.

We have previously found that the enh1-GFP reporter is responsive to multiple Hox genes, including *Drosophila abdB* and *Ubx* and mammalian HoxA9 and HoxB8 (Arya et al, 2015). Furthermore, overexpression of *antp*, *Ubx*, *abdA* and *abdB* have all been shown to ectopically kill embryonic and post-embryonic neural stem cells (Bello et al, 2003; Arya et al, 2015). However, it is unclear what role endogenous expression levels of Antp might play, as Antp is not normally expressed in thoracic neural stem cells (Bello et al, 2003) and our enh1 reporters are not normally active in the thoracic domain (Fig 4; Arya et al, 2015). Additionally, we note that AbdA is endogenously expressed at high levels in many cells that do not die and do not express the enh1 reporters, such as neurons (seen in Fig 2B). These observations suggest that additional factors are involved in determining the context-dependent effects of Antp and AbdA on the activity of the cell death pathway.

### Implications for human disease

Human heterozygous loss of function of Sox2 leads to Anophthalmia-Esophageal-Genital (AEG) syndrome, characterized by failed development or malformation of the eyes, neurocognitive impairments, esophageal-tracheal atresia and urogenital abnormalities (Williamson et al, 2006). In the developing vertebrate gut, Sox2 expression is restricted to anterior tissues and it is highly expressed in the endoderm of the esophagus, trachea and anterior stomach (Ishii et al, 1998; Williamson et al, 2006; Que et al, 2007; Hagey et al, 2018). A similar regional restriction is seen in the *Drosophila* gut, where *Dichaete* is expressed in the posterior gut tissue (Sánchez Soriano & Russell, 2000; Fig S14). In Sox2- hypomorphic mice, the foregut endoderm ectopically expresses the transcription factor and airway marker Nkx2.1, leading to failed separation of the developing esophageal and tracheal tubes, and mis-specification of the esophagus into trachea (Que et al, 2007). This suggests that the role of Sox2 in vertebrate foregut development is to maintain patterning and cell identity. Interestingly, it has been shown that the failure of hindgut development in *D^87^* embryos can be rescued by ectopic expression of *decapentaplegic* (*dpp*), a TGF-beta homologue and marker of the central hindgut (Sánchez Soriano & Russell, 2000). This suggests a conserved role for *Dichaete* in promoting endodermal cell identity, although the expression compartments of *Dichaete* and Sox2 in the developing gut have diverged. We have found that hindgut development is rescued in the *D^87^,MM3* background, indicating that cell death contributes to the hindgut defects upon loss of *Dichaete*. The ectopic expression of *dpp* in the hindgut of *D^87^* embryos may restore cell viability by re-activating endogenous endodermal patterning downstream of *Dichaete*.

Consistent with the failure of proper tissue development upon loss of function of Sox2, high levels of Sox2 are found in many human cancers at both transcript and protein levels. Of particular note is the observation of increased Sox2 expression in samples obtained from glioblastoma patients, and an association of high Sox2 levels with worse prognosis (Ben-Porath et al, 2008; Annovazzi et al, 2011; Alonso et al, 2011; Brennan et al, 2013; de la Rocha et al, 2014; Sathyan et al, 2015; reviewed in Garros-Regulez et al, 2016). Glioblastoma tumors are thought to be seeded and maintained by a subpopulation of glioma stem cells (GSCs), which show enrichment for Sox2 expression and whose continued proliferation and tumor-formation ability depends on Sox2 (Gangemi et al, 2009; Ikushima et al, 2009; Alonso et al, 2011; Hagerstrand et al, 2011; Suva et al, 2014). However, the contribution of cell death to the decreased proliferative state of GSCs upon Sox2 knockdown has not been measured. Gangemi et al (2009) state that no difference in TUNEL staining was observed upon silencing of Sox2 in GSC cultures, in contrast to the observed effects of loss of Sox2 *in vivo* in mice (Favaro et al, 2009). As we noted in the introduction to this study, the role of apoptosis in driving observed phenotypes cannot be conclusively determined without including experimental measures to block cell death. For the first time, we have directly examined the contribution of cell death to phenotypes observed upon loss of *Dichaete*, and we find that multiple developmental defects observed in these mutant embryos can be rescued by preventing apoptosis. In addition, our observation that *Dichaete* overexpression can serve to inhibit activation of the cell death pathway in *Drosophila* offers a mechanism through which Sox2 may promote tumorigenic activity.

## ACKNOWLEDGEMENTS

The authors are grateful to Richa Arya, Seda Gyonjyan and Tatevik Sarkissian for generating the enh1-dsRed and enh1-GFP constructs, as well as to the Bloomington Drosophila Stock Center for fly lines and the Developmental Studies Hybridoma Bank for antibodies. We would particularly like to thank Barbara Nambu and the lab of the late John Nambu for fly stocks and antibodies. This research was supported by the National Institute of Health R01GM110477, R21NS120141 and an MGH Interim support fund to KW.

---

**Figure S1.**
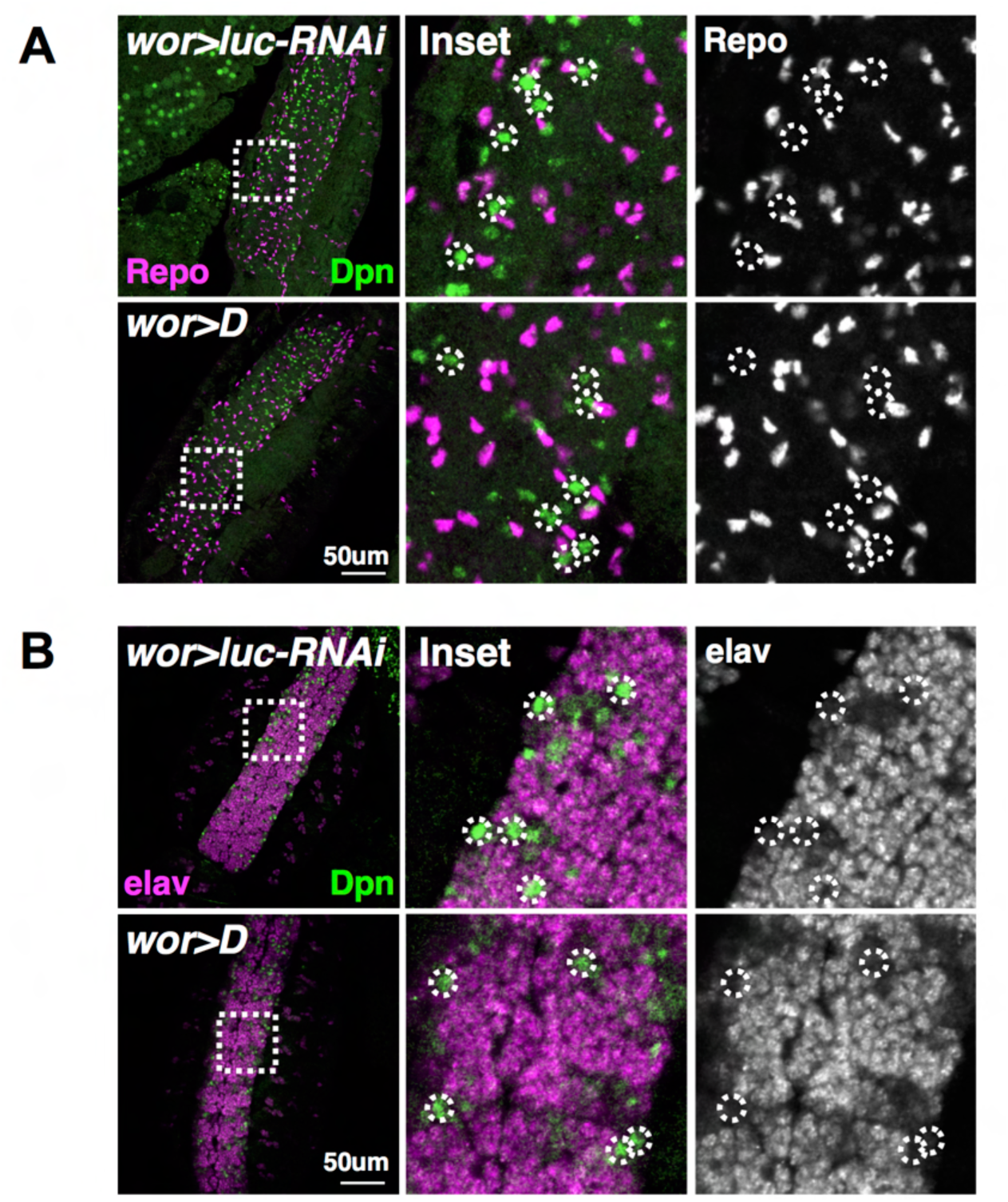
Dichaete-rescued neural stem cells do not express markers of differentiation. **A)** Stage 16 embryos from *wor>dsRed* crossed to *luc-RNAi* or *UAS-D*, stained with anti-Dpn to visualize neural stem cells and anti-Repo to visualize glial nuclei. Images are single confocal slices. **B)** Stage 16 from *wor>dsRed* crossed to *luc-RNAi* or *UAS-D*, stained with anti-Dpn to visualize neural stem cells and anti-elav to visualize neuronal nuclei. Images are single confocal slices.

**Figure S2.**
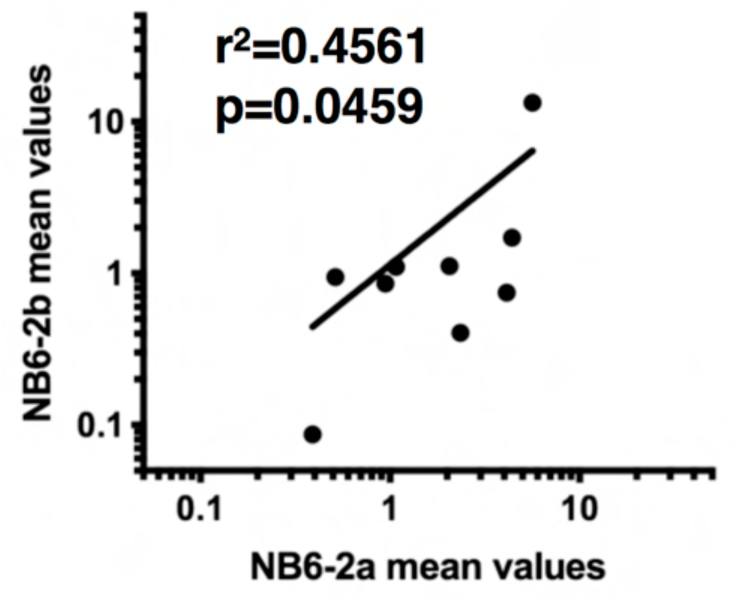
Repeated independent measures of protein expression in NB6-2 show positive correlation. Average intensity measures of AbdA, Grh and D at each developmental stage for NB6-2^a^ (*abdB-GAL4*) and NB6-2^b^ (*calx-GAL4*) were paired and plotted against each other. A Pearson correlation r^2^ value of 0.4561 with a p-value of 0.0459 were obtained, indicating a significant positive relationship between our repeated measures of protein expression in the same neural stem cell across multiple embryos using two independent driver lines.

**Figure S3.**
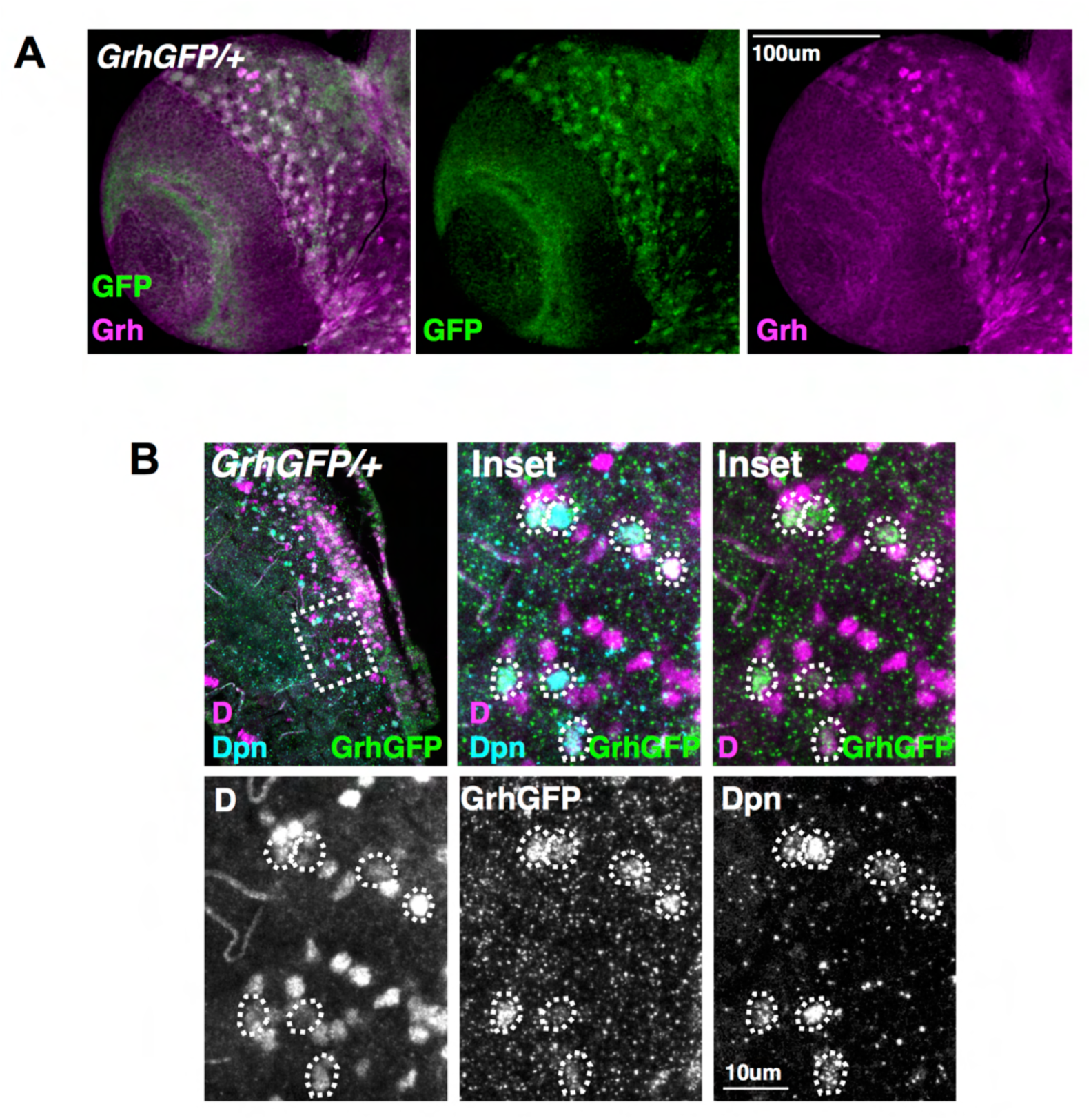
GrhGFP and Dichaete are co-expressed in post-embryonic abdominal neural stem cells. **A)** CNS were dissected from *Grh.GFP/+* wandering L3 larvae and stained with anti-GFP and anti-Grh to determine the extent of overlap between tagged and endogenous proteins in the brain. **B)** Stage 17 *Grh.GFP/+* embryo stained with anti-GFP, anti-D and anti-Dpn to visualize neural stem cells.

**Figure S4.**
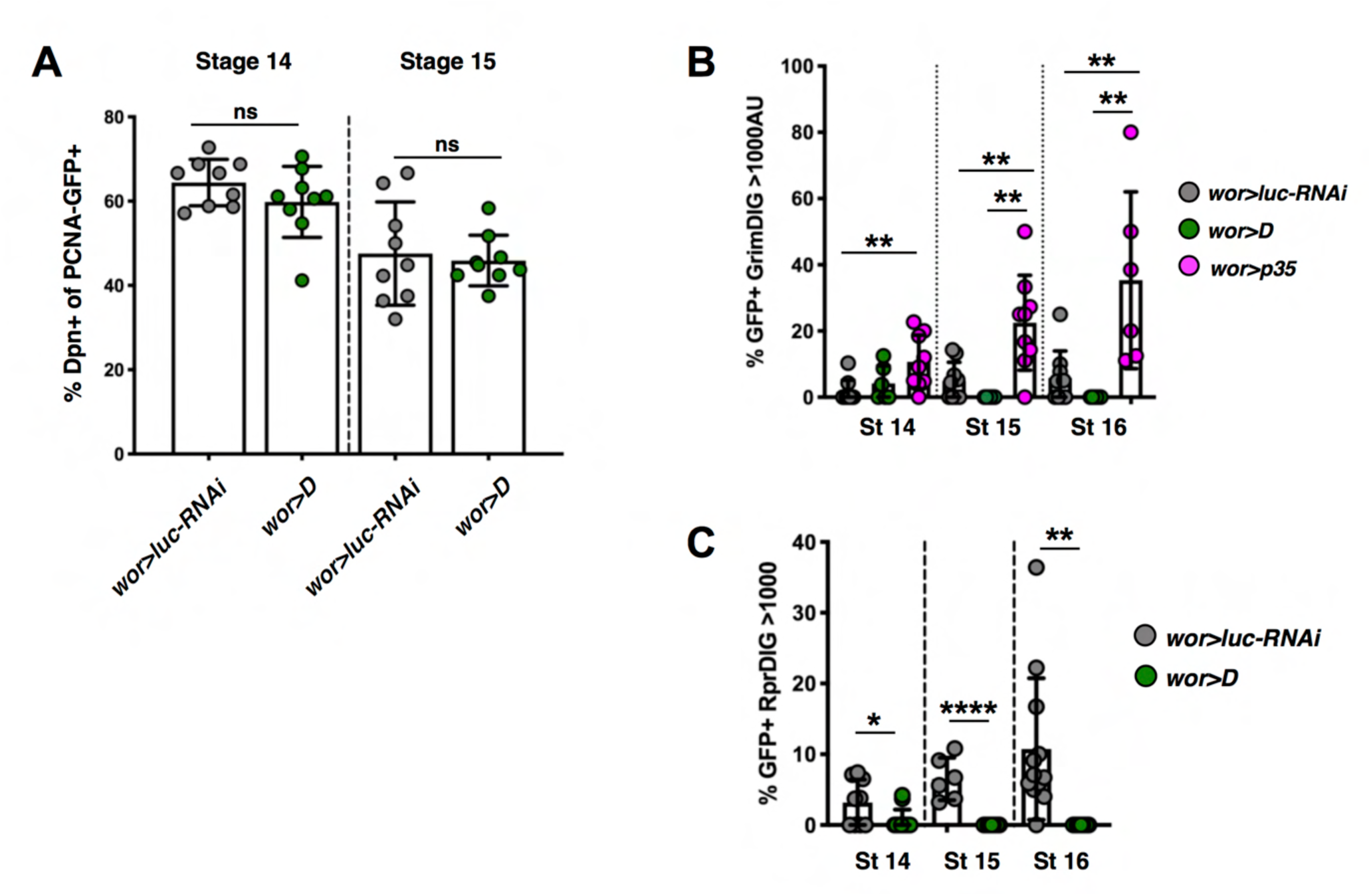
Dichaete downregulates *grim* and *rpr* transcription. **A)** Quantification of Dpn expression in PCNA-GFP+ cells in *wor>luc-RNAi* and *wor>D* embryos at stages 14 & 15. **B)** Quantification of proportion of PCNA-GFP+ cells with GrimDIG signal >1000AU at stages 14-16. This threshold was determined using the qualitative expression assignments shown in Fig 2B&D, as all cells with GrimDIG >1000AU were assigned as having high expression levels. **C)** Quantification of RprDIG signal as in **B**.

**Figure S5.**
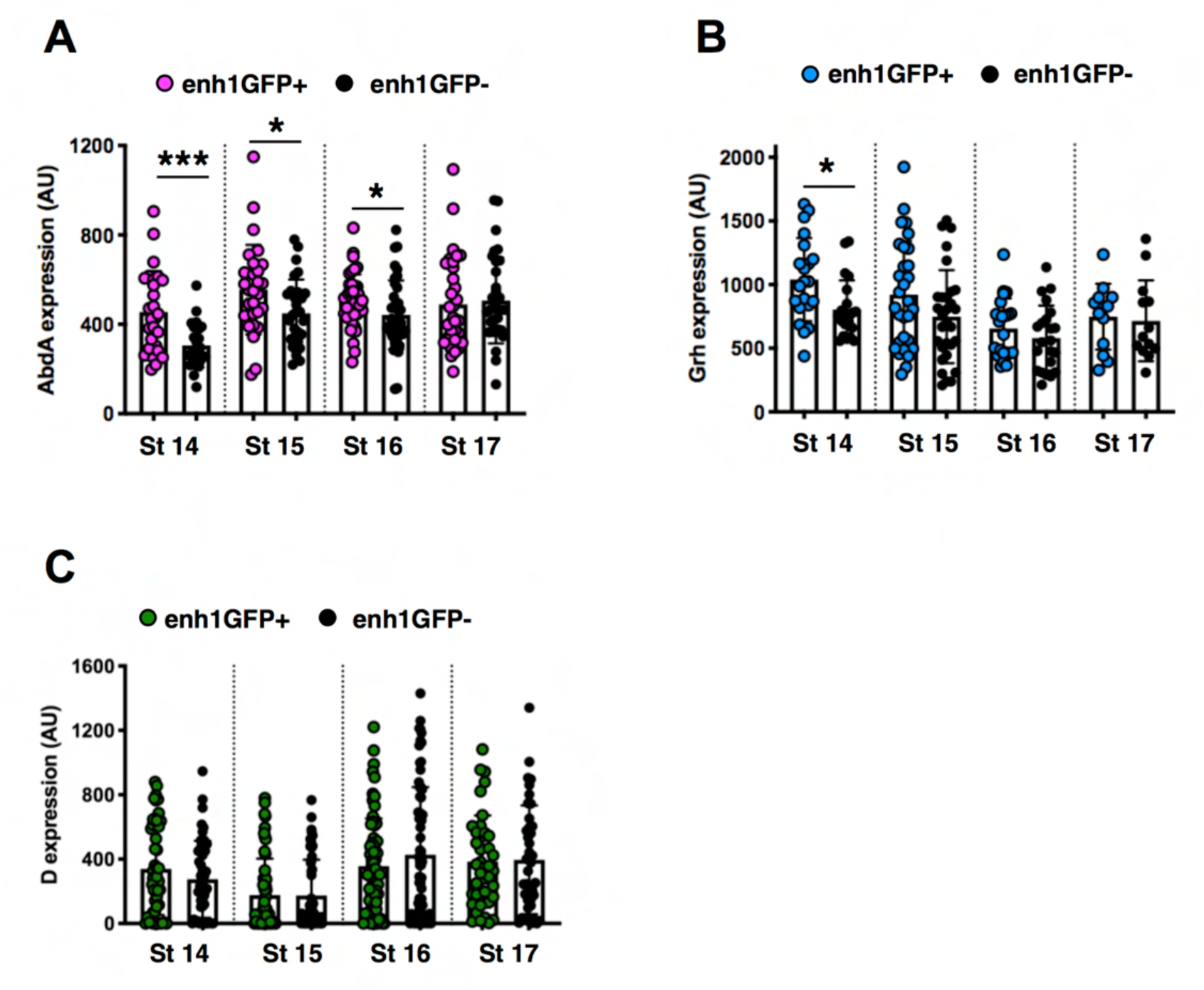
AbdA is expressed at higher levels in enh1-GFP+ neural stem cells compared to enh1-GFP- neural stem cells. **A)** Quantification of AbdA protein expression in Dpn+ cells, categorized as enh1-GFP+ or enh1-GFP-. **B)** Quantification of Grh protein expression, as in **A**. **C)** Quantification of D protein expression, as in **A**.

**Figure S6.**
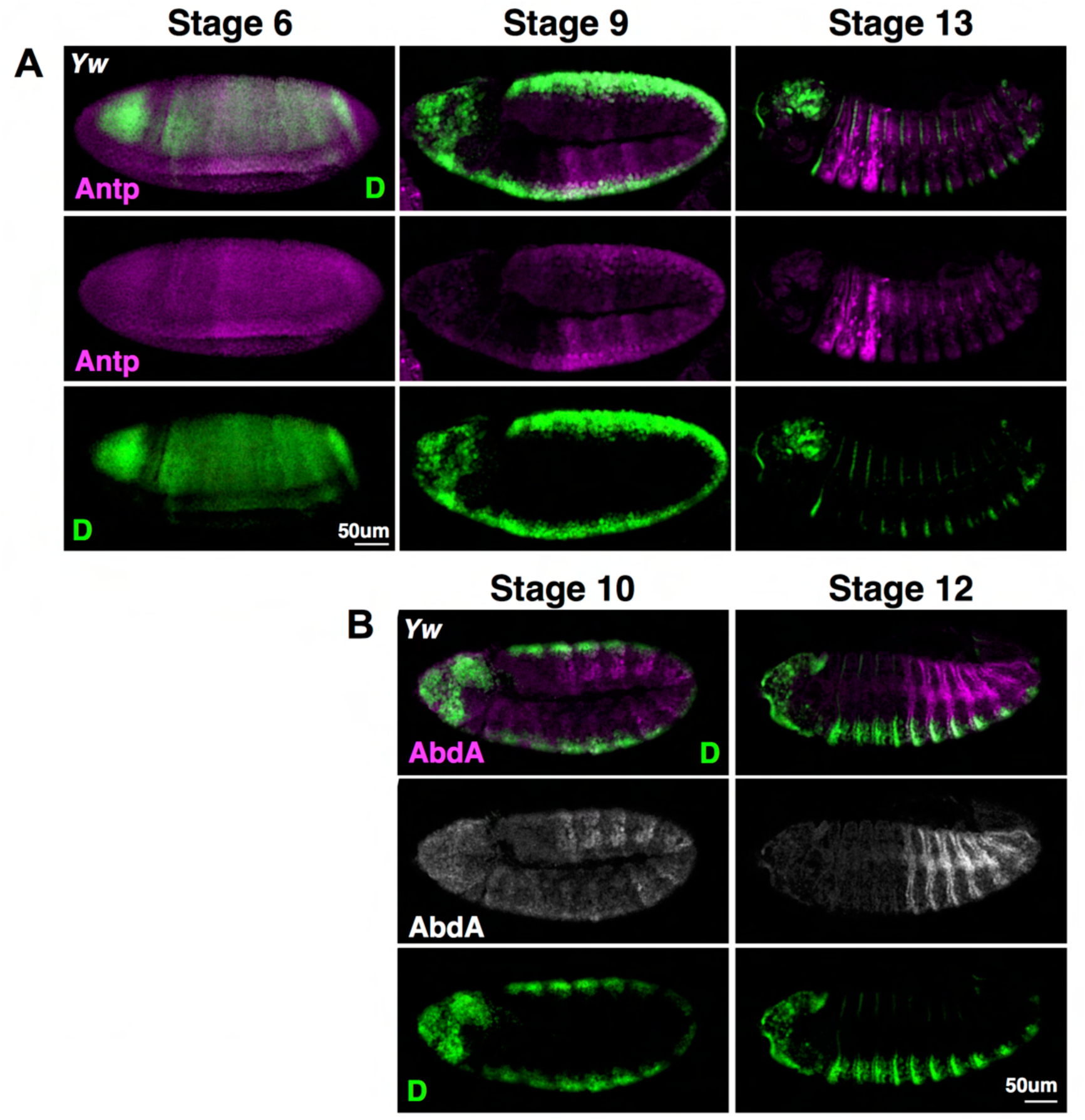
Dichaete is co-expressed with Antp and AbdA in the neuroectoderm in early embryos. **A)** *Yw* embryos were stained with anti-Antp and anti-D, images are single confocal slices. **B)** *Yw* embryos were stained with anti-AbdA and anti-D, images are single confocal slices.

**Figure S7.**
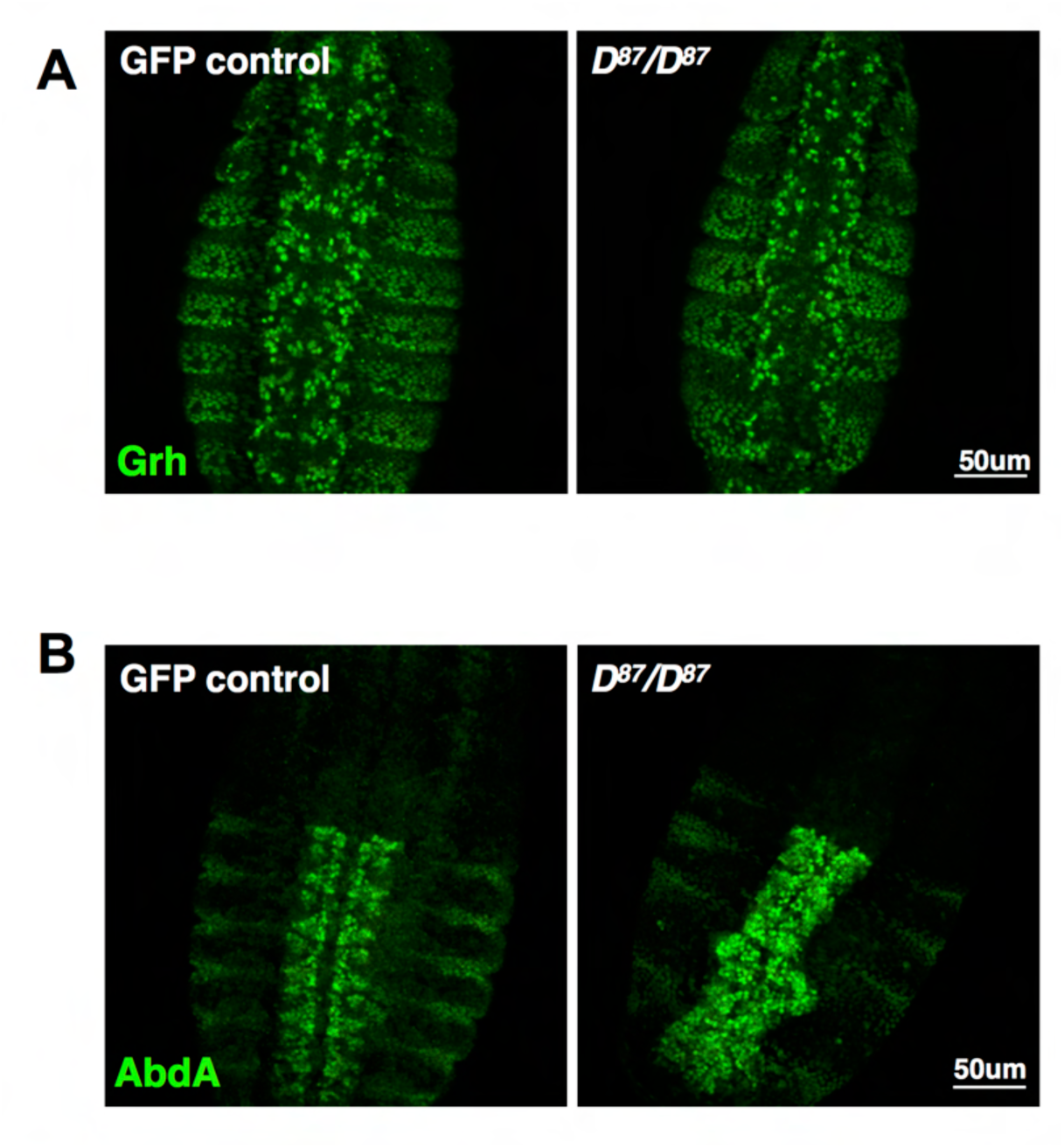
Grh and AbdA are not misregulated in D87 mutant embryos. **A)** Stage 15 embryos from the *D^87^/TM3,GFP* stock stained with anti-Grh. GFP staining (not shown) was used to identify *D^87^/D^87^* mutant embryos. **B)** Stage 15 embryos as above, stained with anti-AbdA. No ectopic expression in the thorax is seen in *D^87^/D^87^* mutant embryos.

**Figure S8.**
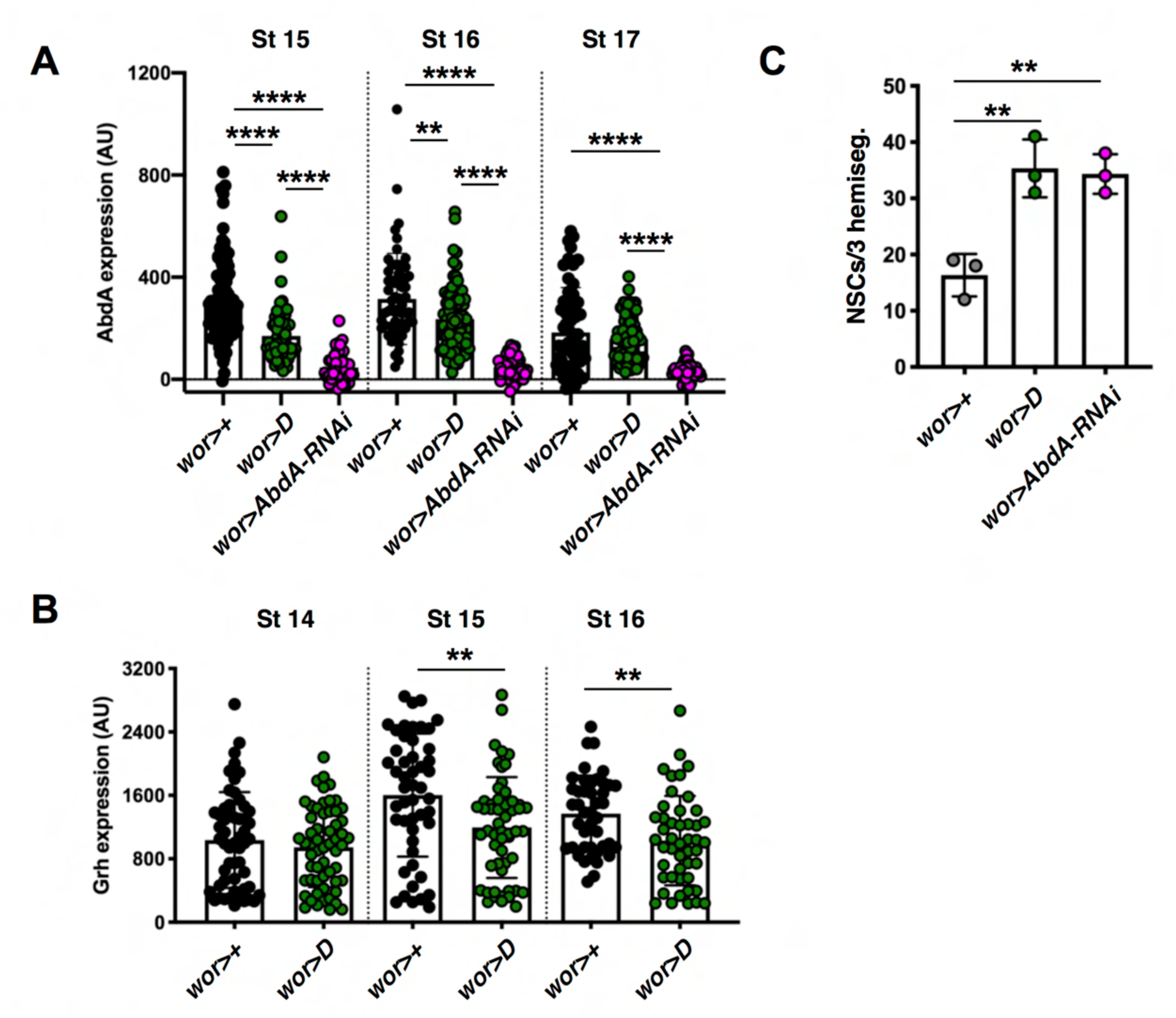
Dichaete overexpression downregulates Grh and AbdA protein expression. **A)** Quantification of AbdA expression in Dpn+ cells in embryos from *wor>dsRed* crossed to the indicated genotypes. **B)** Quantification of Grh expression in Dpn+ cells in *wor>luc-RNAi* and *wor>D* embryos. **C)** Quantification of abdominal neural stem cell survival at stage 17 in *wor>luc-RNAi, wor>D* and *wor>AbdA-RNAi* embryos.

**Figure S9.**
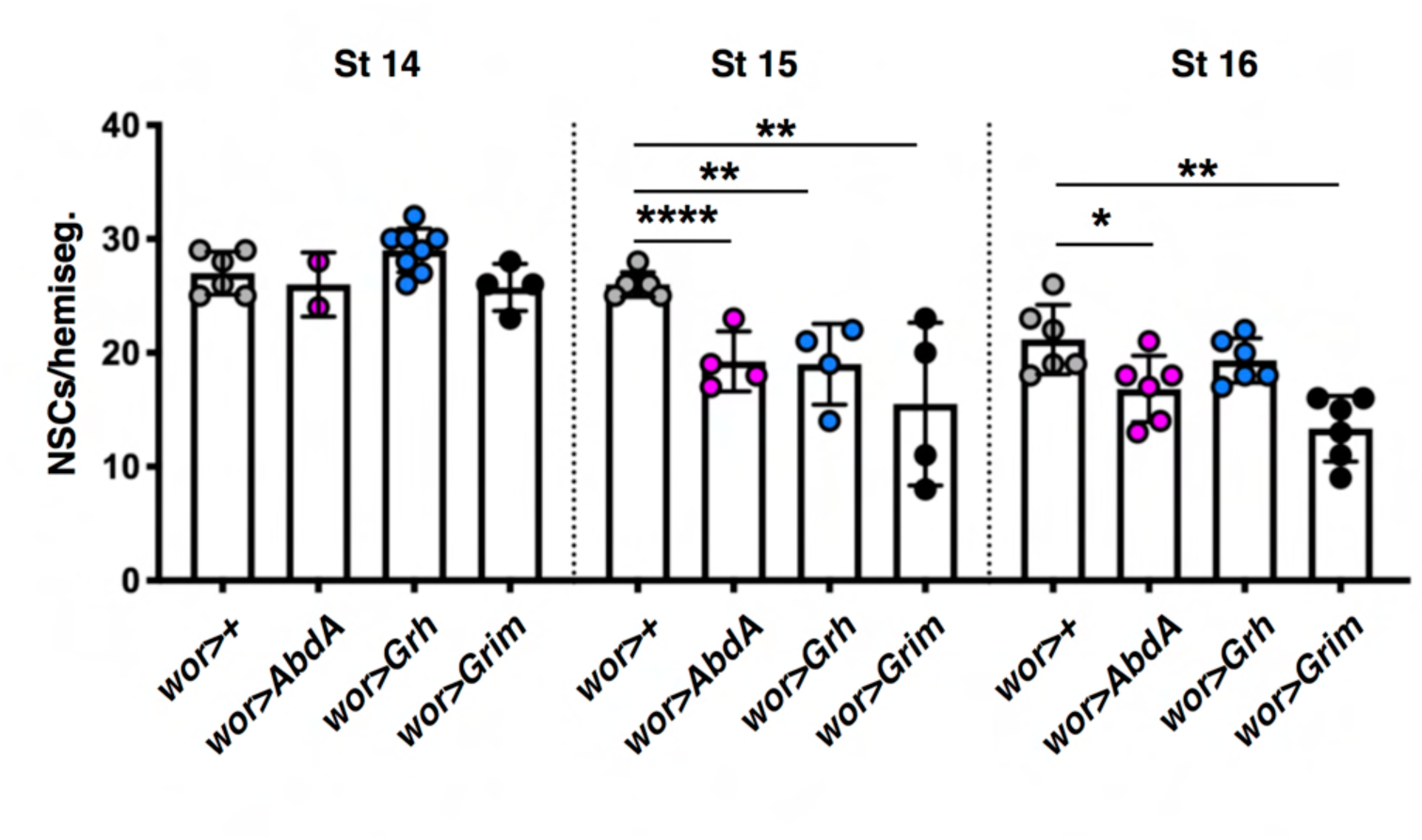
AbdA and Grh are sufficient to kill thoracic neural stem cells. Quantification of neural stem cell survival in 2 thoracic hemisegments in embryos from *wor>dsRed* crossed to the indicated genotypes. *UAS-Grim* is included as a positive control.

**Figure S10.**
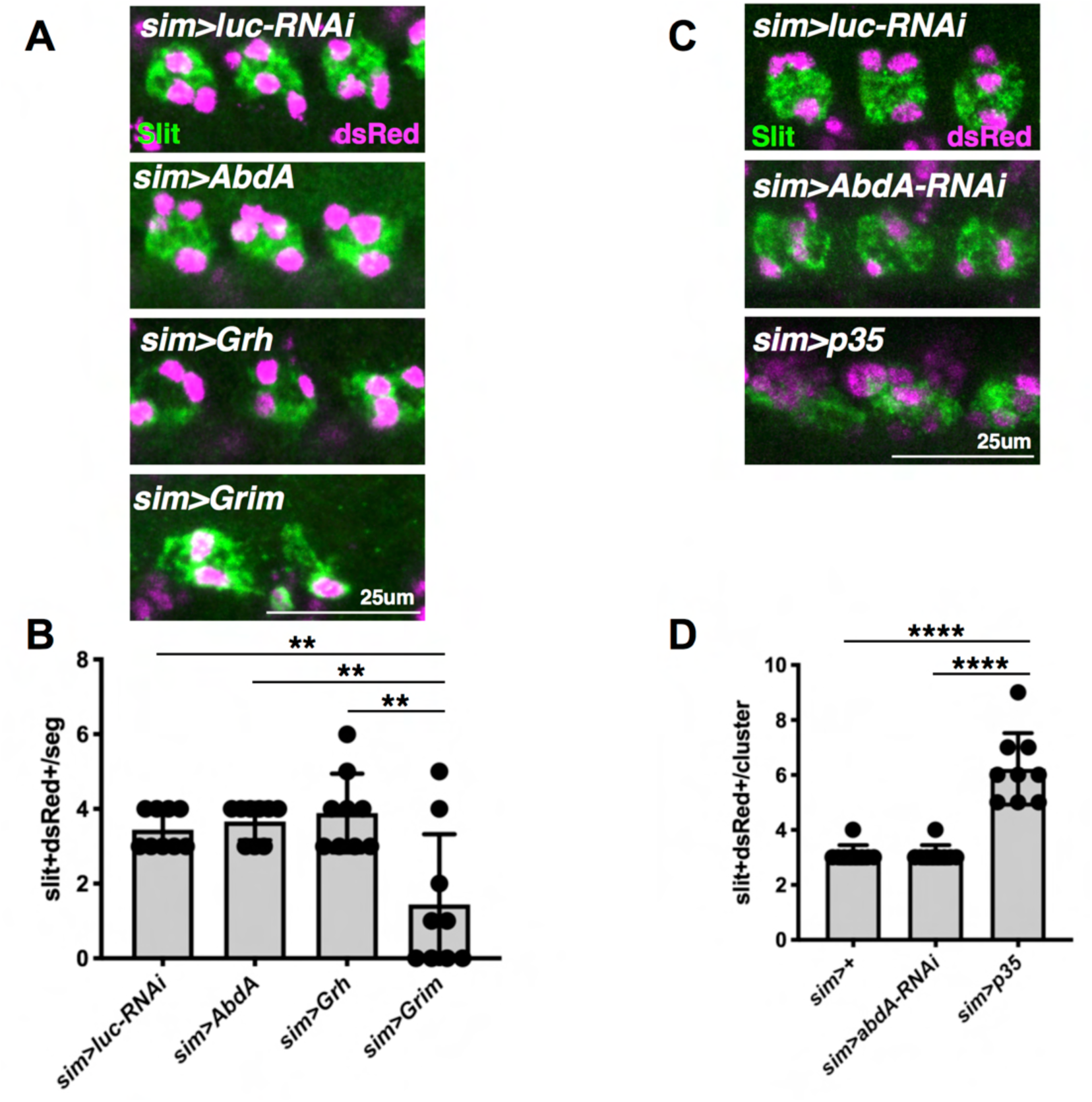
AbdA and Grh do not kill midline glia, and AbdA is not required for their death. **A)** Stage 17 embryos from *sim>dsRed* crossed to the indicated genotypes, stained with anti-Slit. Images are maximum projections through the Slit signal. **B)** Quantification of Slit+dsRed+ nuclei per midline cluster, shown in **A**. n=3 clusters from 3 embryos per genotype. **C)** Stage 17 embryos from *sim>dsRed* crossed to the indicated genotypes, stained with anti-Slit. Images are maximum projections through the Slit signal. **D)** Quantification of Slit+dsRed+ nuclei per midline cluster, shown in **B**. n=3 clusters from 3 embryos per genotype.

**Figure S11.**
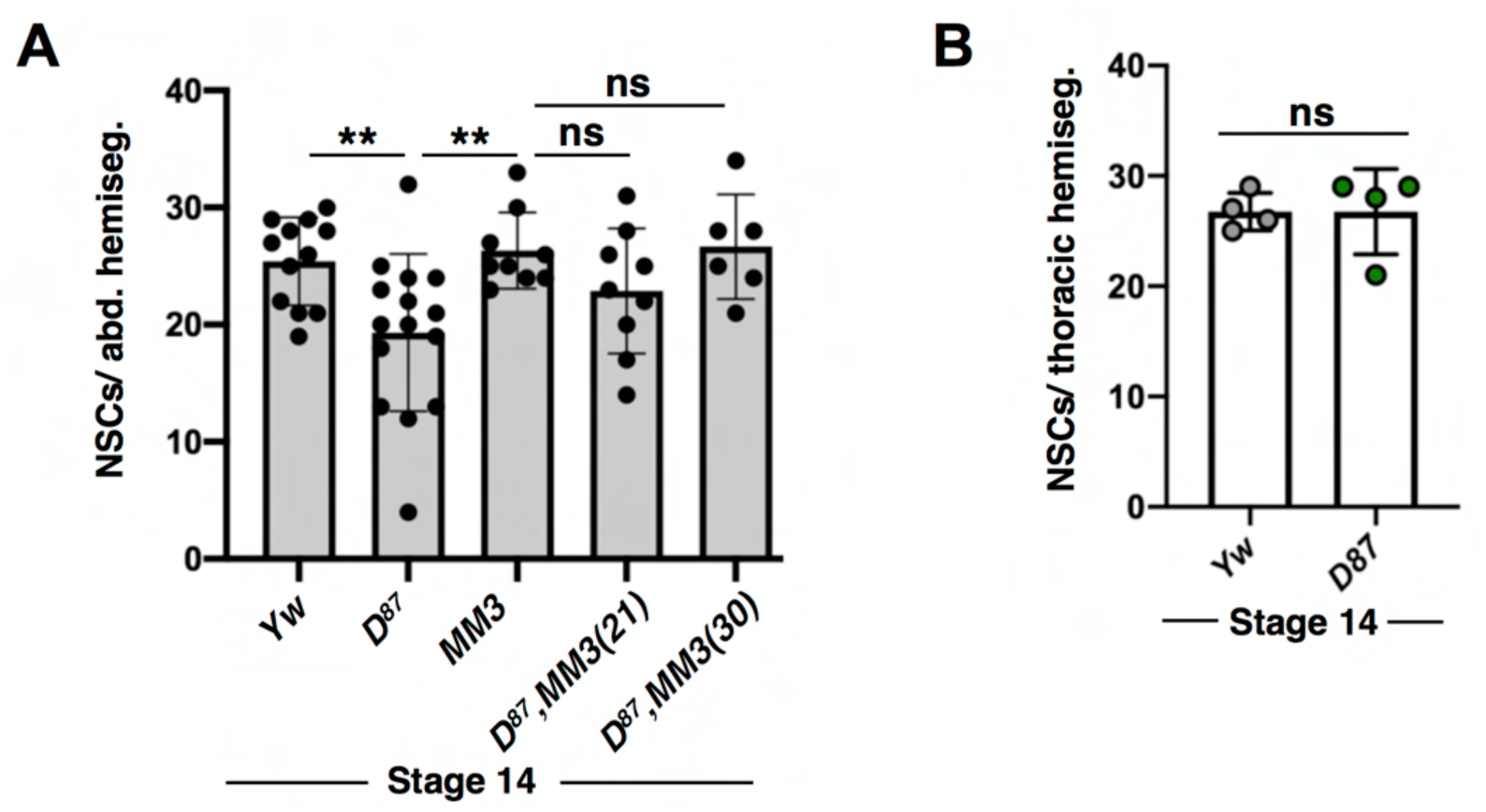
Early abdominal neural stem cell loss in *D^87^* mutants is rescued by MM3. **A)** Quantification of abdominal neural stem cell numbers at stage 14, determined by anti-Dpn and anti-AbdA staining. n=3 hemisegments from ≥2 embryos per genotype. **B)** Quantification of thoracic neural stem cell numbers at stage 14, determined by anti-Dpn and anti-AbdA staining. n=2 hemisegments from 2 embryos per genotype.

**Figure S12.**
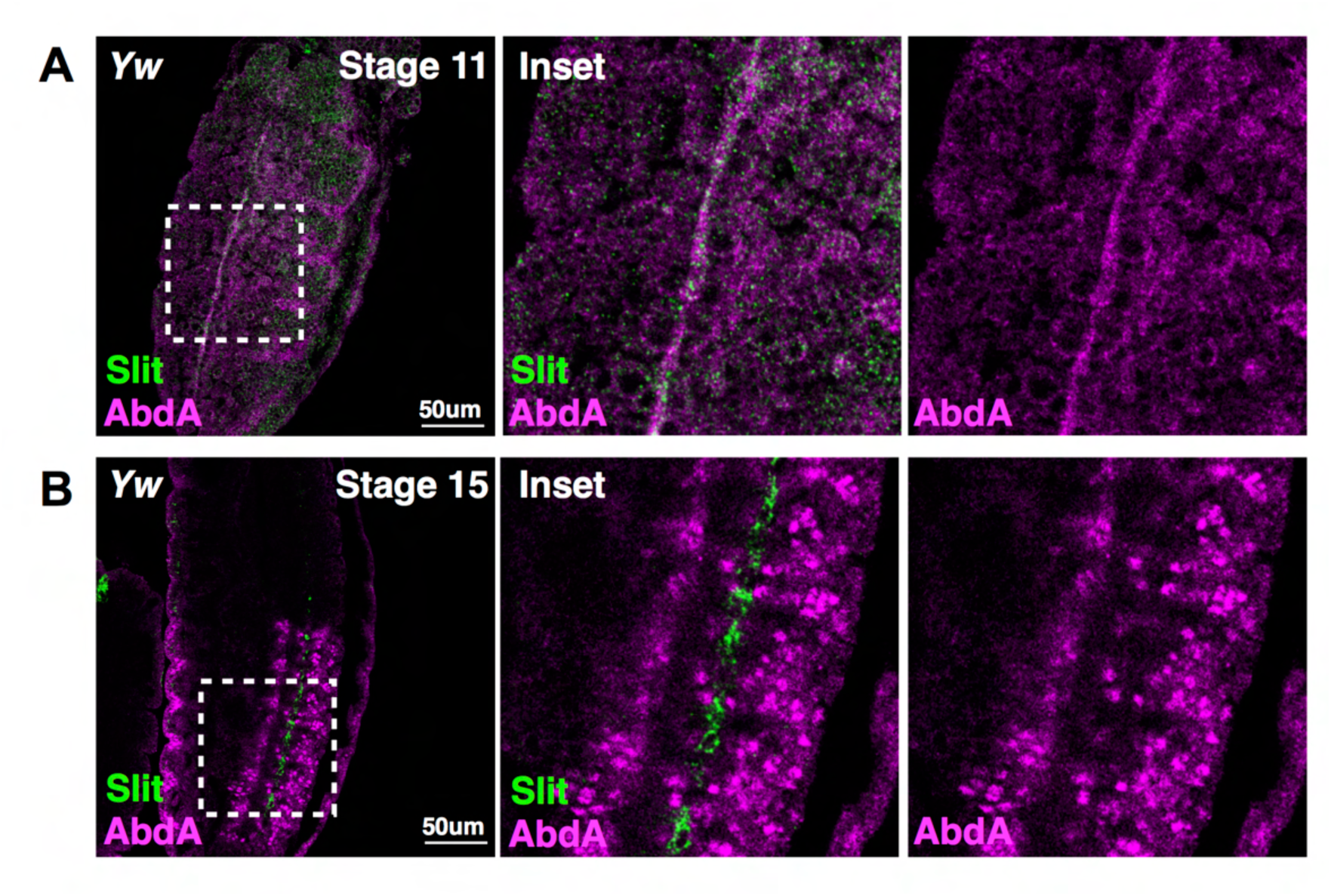
AbdA is expressed along the midline of early embryos. **A)** Stage 11 *Yw* embryo stained with anti-Slit and anti-AbdA, image is a single confocal slice. **B)** Stage 15 *Yw* embryo, as in **A**.

**Figure S13.**
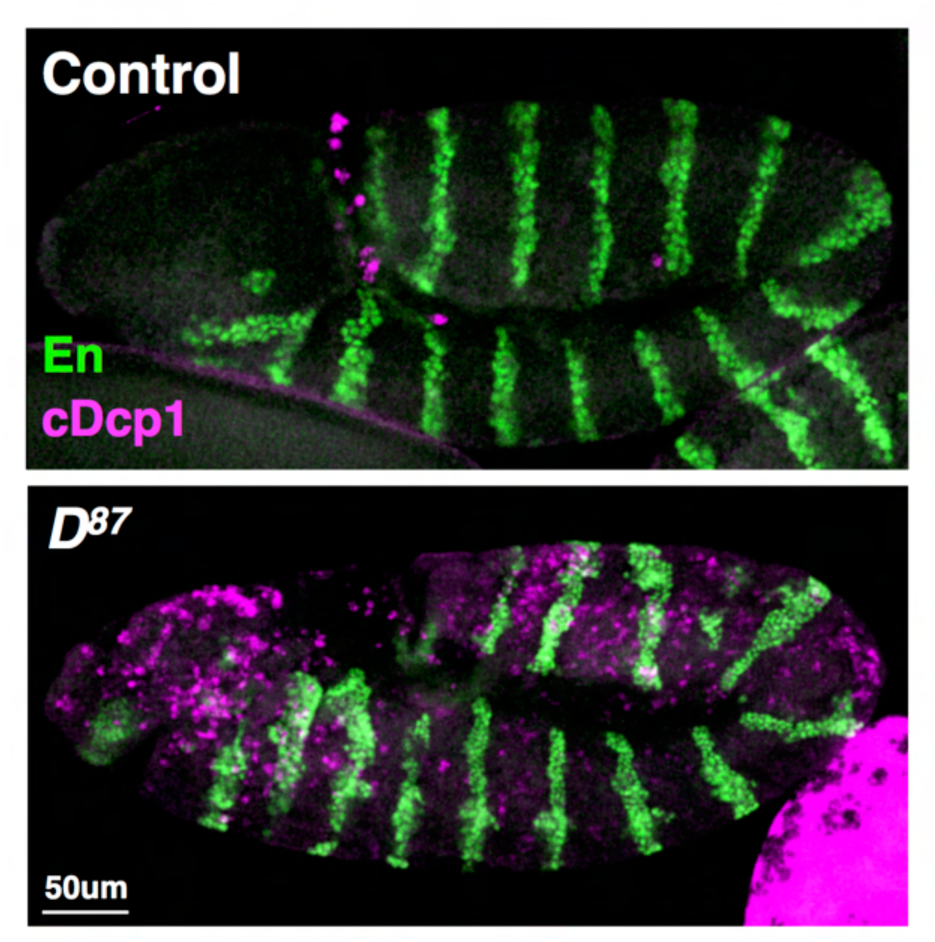
Ectopic caspase activation in *D^87^* mutant embryos. **A)** GFP+ control embryo at stage 10/11 stained with anti-En and anti-cDcp1, image is a maximum projection. **B)** *D^87^* mutant embryo at stage 10/11, as in **A**.

**Figure S14.**
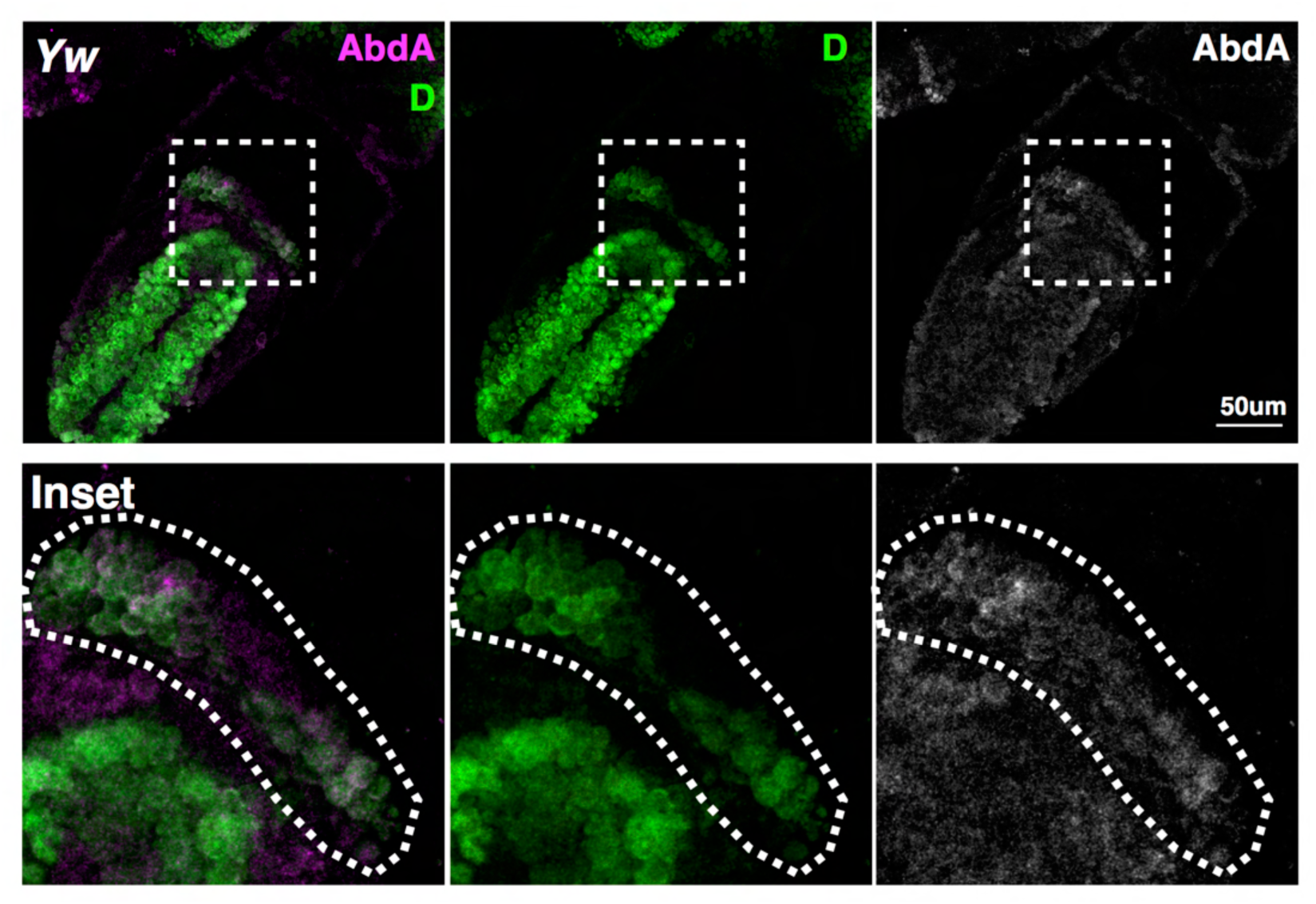
Dichaete and AbdA are co-expressed in the hindgut primordium. Stage 10 *Yw* embryo stained with anti-D and anti-AbdA, the hindgut primordium is marked by high Dichaete expression and outlined with a dashed line. Image is a single confocal slice.

